# On the interpretation of the operation of natural selection in class-structured populations

**DOI:** 10.1101/2022.05.31.494171

**Authors:** Tadeas Priklopil, Laurent Lehmann

**Affiliations:** Department of Ecology and Evolution, University of Lausanne, Lausanne, Switzerland; Institute of Integrative Biology & Institute of Molecular Plant Biology, ETH Zurich, Zürich, Switzerland

## Abstract

Biological adaptation is the outcome of allele-frequency change by natural selection. At the same time, populations are usually class structured as individuals occupy different states such as age, sex, or stage. This is known to result in the differential transmission of alleles through non-heritable fitness differences called class transmission, also affecting allele-frequency change even in the absence of selection. How does one then isolate the allele-frequency change due to selection from that owing to class transmission? We decompose one-generational allele-frequency change in terms of effects of selection and class transmission, and show how reproductive values can be used to reach a decomposition between any two distant generations of the evolutionary process. This provides a missing relationship between multigenerational allele-frequency change and the operation of selection. It also allows to define a measure of fitness summarizing the effect of selection in a multigenerational evolutionary process, which connects asymptotically to invasion fitness.

## Introduction

Consider the following situation you may encounter as an evolutionary biologist. Suppose you observe a population between two distant points in time, and empirically measure at each census point the total number of individuals in the population and the alleles they carry at some locus of interest. Suppose further you can also measure the survival and number of descendants of these individuals, and that these measurements can be subdivided into different reproductive classes the individuals carrying the alleles can be in, such as sex, age, stage or habitat. The overall time of observation is moreover assumed short enough to neglect the appearance of new mutations and the population is closed and large enough to neglect the effect of migration and genetic drift. Then, any change in allele frequency must have occurred due to natural selection and/or the process of class transmission, where the latter is the differential reproductive success of an individual owing to differential classes in which individuals reproduce and survive. This is also known to alter allele frequencies in class-structured populations even in the absence of selection [e.g., Charlesworth, 1994, Crow, 1979, Grafen, 2006, Leturque and Rousset, 2002, Stubblefield and Seger, 1990, Taylor, 1990]. However, only natural selection can cause allelic fixations under the above scenario and is the only evolutionary force that shapes biological adaptations [Barton et al., 2007, Fisher, 1930, Futuyama, 2017, Lynch and Walsh, 2018]. It is therefore natural to ask how much of the measured change in allele frequency is caused by natural selection and how much by class transmission, and how much of the overall change can be attributed to each demographic time step of the evolutionary process? To answer this question requires a careful account of the operation of natural selection and class transmission in class-structured populations.

Most previous theoretical work on evolution in class-structured populations, however, has focused on biological scenarios where class transmission has a negligible effect on the evolutionary dynamics [e.g., Barfield et al., 2011, Caswell, 2001, Engen et al., 2014, Gardner, 2015, Grafen, 2006, 2015, Leturque and Rousset, 2002, Lion, 2018a, Lion and Gandon, 2022, Priklopil and Lehmann, 2020, Rousset, 2004, Taylor, 1990]. This is the case, for instance, in a large-time limit when either survival and reproduction is frequency- and density-independent [Engen et al., 2014, Grafen, 2006, 2015] or when selection is assumed weak [Barfield et al., 2011, Leturque and Rousset, 2002, Lion, 2018a, Lion and Gandon, 2022, Priklopil and Lehmann, 2020, Rousset, 2004, Rousset and Ronce, 2004, Taylor and Frank, 1996]. In this work, the asymptotic evolutionary success of an allele is characterised in terms of a weighted allele frequency (or weighted trait average) with weights as reproductive values, which can loosely be interpreted as genetic contributions of individuals to the future of the population [e.g., Barton and Etheridge, 2011]. The main justification that has been given for using reproductive values as weights is that the long-term reproductive output of offspring produced into different classes is converted to a common measure that can be compared and added up [Taylor and Frank, 1996], and as a consequence the dynamics of weighted averages behave ‘nicely’ in that they cancel any allele-frequency change caused by non-heritable fitness differ-ences [Crow, 1979, Grafen, 2006]. The resulting dynamics governed purely by reproductive-value weighted fitness differentials is then taken as a measure of selection [Engen et al., 2014, Grafen, 2006, 2015, Leturque and Rousset, 2002, Lion, 2018a, Lion and Gandon, 2022, Priklopil and Lehmann, 2020, Rousset, 2004, Rousset and Ronce, 2004, Taylor and Frank, 1996]. However, in this previous work it is not clear how the reproductive-value weighted frequency connects to the standard arithmetic average allele frequency describing the unfolding of the evolutionary process. As a consequence, no explicit formal nor verbal biological reason has been given in why reproductive-value weighted allele-frequency is an appropriate measure of selection, and how to interpret it in terms of the arithmetic allele frequency. Further, for non-asymptotic finite-time evolutionary processes, class transmission does contribute to allele-frequency change and hence cannot be ironed away by using reproductive values as weights.

The aim of this paper is to provide an operational decomposition of the contribution of natural selection and that of class transmission to the arithmetic average allele-frequency change in a class-structured population, where individuals may experience frequency- and density-dependent interactions as well as environmental fluctuations. In this decomposition, we consider a change over an arbitrary number of demographic time steps and allocate fractions of the average allele-frequency change to each step of this evolutionary process. Our analysis shows, in particular, that the previously used reproductive-value weighted allele-frequency change is exactly the arithmetic allele-frequency change caused by selection when ‘assessed’ at the end of the observed evolutionary process. We also provide the expression for the previously unaccounted allele-frequency change due to class transmission, such that the sum of these two contributions gives the total arithmetic allele-frequency change attributed to any given generation but assessed at the end of the observed process. These calculations highlight that the allele-frequency change caused by natural selection and class transmission during any demographic time step is tied to the entire evolutionary process of interest and hence cannot be studied in isolation. Finally, we provide a detailed and biologically explicit definition of reproductive value, and provide a representation of fitness summarizing the effect of selection and class transmission in the multiple time-step evolutionary process.

Our analysis proceeds in two main steps. In order to motivate the formalization of allele-frequency change over an evolutionary process spanning multiple demographic time steps, we first consider the traditional approach in calculating the allele-frequency change over a single time-step. Second, we turn to the multiple time-step evolutionary process, which forces us to reconsider some of the intuition gained from the single time-step change. Here, we derive our main results, discuss the related literature, and then connect our results to earlier work on asymptotic evolutionary dynamics.

### Single Generational Process in Class-Structured Populations

#### The Model

For simplicity but without loss of generality, we consider a population of haploid individuals that reproduce asexually, each of which is characterized by one of a finite number of alleles that segregate in the population. The population is assumed class-structured so that each individual belongs to one of a finite number *n*_*C*_ of classes (age, stage etc.) and can possibly transition between the classes, e.g., via survival, maturation or physiological development. We furthermore assume that reproduction and class transition occurs at discrete demographic time steps, that there are no mutations, and that the population is closed and large enough to ignore the effect of migration and genetic drift. Then, the state of the population can be fully characterized by the vector **n**_*t*_ collecting elements of the form *n*_*t*_(*a, i*), which stands for the number of individuals in class *a* with allele *i* at time *t*. This number satisfies the recursion

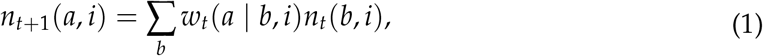

which defines a multi-type population process where we use the shorthand notation *w*_*t*_(*a* | *b, i*) = *w*_*t*_(*a* | *b, i*, **n**_*t*_) to represent the class-specific individual fitness of a carrier of allele *i*. The class-specific individual fitness *w*_*t*_(*a* | *b, i*) is defined as the expected number of settled ‘offspring’ in class *a* with allele *i* at *t* + 1 produced by a single ‘parent’ with allele *i* in class *b* at *t* including the surviving self (hence quotation marks). This thus gives the number of gene copies *a* at time *t* + 1 descending from a parent *b* carrying allele *i* and alive at *t*. Because the class-specific individual fitness may depend in an arbitrary way on the population state **n**_*t*_, the model allows for arbitrarily complicated frequency- and density-dependent interactions between individuals. Because this individual fitness is also indexed by *t* it further allows for arbitrary extrinsic environmental fluctuations affecting reproduction and class transition. We sum over all classes in eq. (1) because parents in any class may possibly produce offspring of any other class, and we note that all equations throughout hold for all segregating alleles.

#### Allele-Frequency Change

To analyze allele-frequency change induced by the population process in eq. (1), we need to introduce the following notations (see Table 1 for a summary of all notations). First, we denote by *n*_*t*_(*i*) = ∑_*a*_ *n*_*t*_(*a, i*) the total number of individuals carrying allele *i*, by *n*_*t*_(*a*) = ∑_*i*_ *n*_*t*_(*a, i*) the total number of individuals in class *a* and by *n*_*t*_ = ∑_*a,i*_ *n*_*t*_(*a, i*) the total number of individuals in the population. From these quantities, *p*_*t*_(*i*) = *n*_*t*_(*i*)/*n*_*t*_ is the frequency of allele *i, p*_*t*_(*a*) = *n*_*t*_(*a*)/*n*_*t*_ is the frequency of class *a* individuals, *p*_*t*_(*a* | *i*) = *n*_*t*_(*a, i*)/*n*_*t*_(*i*) is the frequency of class *a* individuals amongst those with allele *i* and *p*_*t*_(*i* | *a*) = *n*_*t*_(*a, i*)/*n*_*t*_(*a*) is the frequency of allele *i* within class *a*. From this, and summing eq. (1) over all classes and alleles, we have that 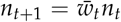, where 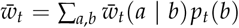 is the average fitness in the population, which is here expressed in terms of the class-specific average fitness 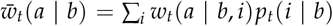 over alleles, while the average fitness of allele *i* itself is 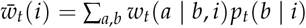.

**Table 1:** Definitions of the variables and main quantities. See main text for further explanations.

| Symbol | Definition |
| --- | --- |
| $n_t(a, i)$ | Number of individuals in class $a$ carrying allele $i$ at time $t$ |
| $n_t(i) = \sum_a n_t(a, i)$ | Total number of individuals carrying allele $i$ at time $t$ |
| $n_t(a) = \sum_i n_t(a, i)$ | Total number of individuals in class $a$ at time $t$ |
| $p_t(i) = n_t(i)/n_t$ | Frequency of individuals in the population carrying allele $i$ at time $t$ |
| $p_t(a \vert i) = n_t(a, i)/n_t(i)$ | Frequency of individuals in class $a$ amongst all the individuals that carry allele $i$ in the population at time $t$ |
| $p_t(i \vert a) = n_t(a, i)/n_t(a)$ | Frequency of individuals carrying allele $i$ amongst all the individuals in class $a$ at time $t$ |
| $w_t(a \vert b, i)$ | Expected number of offspring of class $a$ produced over the time step $[t, t + 1]$ by a single parent in class $b$ carrying allele $i$ |
| $\bar{w}_t(a \vert b) = \sum_i w_t(a \vert b, i)p_t(i \vert b)$ | Expected number of offspring of class $a$ produced over the time step $[t, t + 1]$ by a single average parent in class $b$ |
| $\bar{w}_t(i) = \sum_{a,b} w_t(a \vert b, i)p_t(b \vert i)$ | Expected number of offspring produced over the time step $[t, t + 1]$ by a single average parent $i$ in the population |
| $\bar{w}_t = \sum_{a,b} \bar{w}_t(a \vert b)p_t(b)$ | Expected number of offspring produced over the time step $[t, t + 1]$ by a single average parent in the population |
| $v_t(a)$ | Reproductive value of an individual $a$ at time $t$ in a multigenerational process over $\mathcal{T}$, defined as the expected number of individuals at time $t_f$ that descend under the neutralized process from that individual |
| $\bar{v}_t(i) = \sum_a v_t(a)p_t(a \vert i)$ | Reproductive value of an average $i$ at time $t$ in a multigenerational process over $\mathcal{T}$ |
| $\bar{v}_t = \sum_i \bar{v}_t(i)p_t(i)$ | Reproductive value of an average individual in the population at time $t$ in a multigenerational process over $\mathcal{T}$ |
| $\bar{w}_t(i) = \sum_{a,b} \bar{w}_t(a \vert b)p_t(b \vert i)$ | Expected number of offspring produced over the time step $[t, t + 1]$ by a single $i$ at time $t$ if it were to survive and reproduce as an average individual in the class |
| $n_t^\circ(a, i)$ | Total number of individuals $i$ in class $a$ at time $t$ in the neutralized process where all individuals between $[t_0, t]$ survive and reproduce as average individuals of their class |
| $n_t^\circ(i)$ | Total number of individuals $i$ at time $t$ in the neutralized process where all individuals between $[t_0, t]$ survive and reproduce as average individuals in the class |
| $p_t^\circ(a \vert i)$ | Frequency of $a$ at $t$ amongst all the individuals that carry allele $i$ in the neutralized process that started at time $t_0$ |
| $\bar{w}_t^\circ(i) = \sum_{a,b} \bar{w}_t(a \vert b)p_t^\circ(b \vert i)$ | Expected number of offspring produced over the time step $[t, t + 1]$ by a single $i$ if it were to reproduce as an average individual in the class and that comes from a lineage of individuals that descend under the neutralized process between $[t_0, t]$ |
| $\bar{w}_t^v(i) = \sum_{a,b} v_t(a)w_t(a \vert b, i)p_{t+1}(b \vert i)$ | Expected number of individuals at time $t_f$ that descend under the neutralized process from all offspring produced during $[t, t + 1]$ by a single $i$ carrier |
| $r^\circ(i) = n_t^\circ(i)/n_t(i)$ | Fraction of $i$ at $t$ that come from a lineage of individuals that underwent only the neutralized process during $[t_0, t]$ |
| $n_{\mathcal{C}}$ | Total number of classes in the class-structured population |
| $q_t(a)$ | Arbitrary weight of individuals in class $a$ at time $t$ |
| $q_t(i) = \sum_a q_t(a)p_t(a \vert i)$ | Arbitrary weight of an average individual $i$ in the population at time $t$ |
| $\bar{w}^q(i) = \sum_{a,b} q_{t+1}(a)w_t(a \vert b, i)p_{t+1}(b \vert i)$ | Expected number of offspring produced during $[t, t + 1]$ by a single $i$ when each offspring in each class is weighted by an arbitrary weight $q_{t+1}(a)$ |

The change in frequency Δ*p*_*t*_(*i*) = *p*_*t*+1_(*i*) − *p*_*t*_(*i*) of allele *i* over a single time step [*t, t* + 1], which we refer to as ‘generation’ *t*, is then

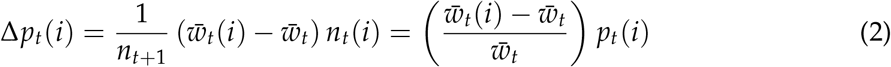

(Appendix A1.1). The first equality makes the cause of allele-frequency change in generation *t* explicit: 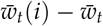 gives the additional number of offspring produced by a single allele *i* carrier compared to an average individual from the total population. Multiplying this with the total number of parents *i* at *t* and dividing by the total number of offspring at *t* + 1 we obtain the change in allele-frequency *i* in generation *t*. The second equality in eq. (2) is the standard representation of allele-frequency change in the absence of mutation and genetic-drift (e.g., Gillespie, 1991, eq. 4.1, Nagylaki, 1992, eq. 2.8 for models without class structure). Indeed, an allele can only increase in frequency if it propagates through the population faster than alleles do on average: allele *i* increases in frequency in generation *t* whenever 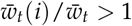.

While it would be tempting to conclude that natural selection is responsible for all the allele-frequency change in eq. (2), as is the case in populations without class structure (Figure 1; Gillespie, 1991, Nagylaki, 1992), this is not true in class-structured populations. Indeed, one can see this by considering a situation where every allele in each class has the same class-specific fitness *w*_*t*_(*a* | *b, i*) = *w*_*t*_(*a* | *b*) for all *i* segregating in the population at all times. This thus defines a neutral (population) process whereby eq. (2) reduces to

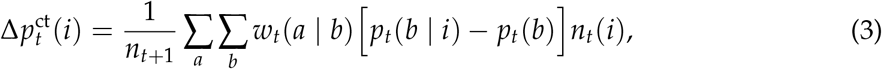

where we used the superscript ‘ct’ to indicate that all change in allele frequency 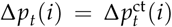 is caused by class transmission only (Appendix A1.1). Equation (3) shows that non-selective allele-frequency change occurs due to a non-uniform distribution of alleles across classes and proceeds until all alleles are fully transmitted through all the classes resulting in *p*_*t*_(*b* | *i*) = *p*_*t*_(*b*), a point at which there is zero allele-frequency change.

**Figure 1:**
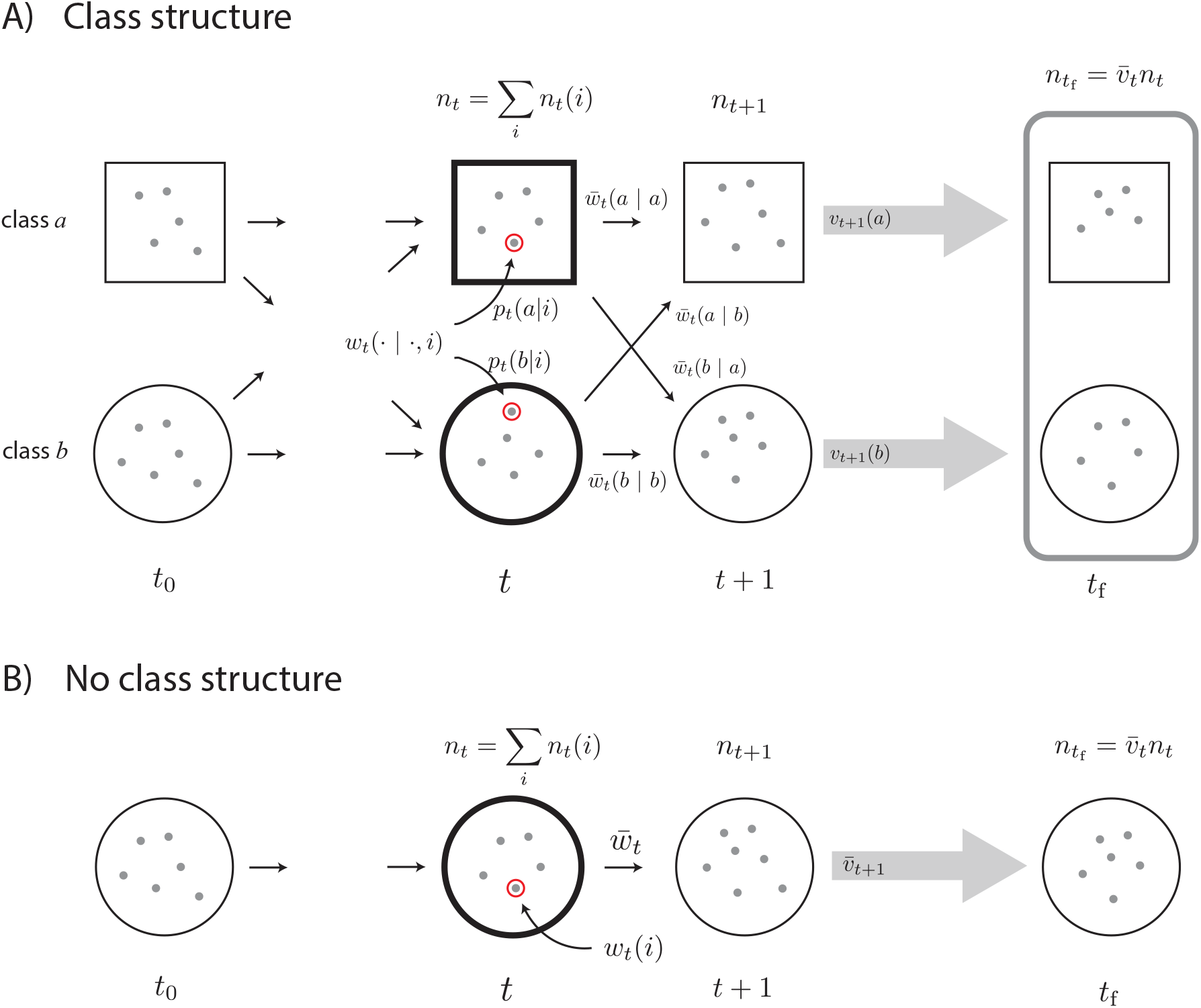
**Panel A)** Evolutionary dynamics in a class-structured population, where at each point in time *t* the population is separated into two classes *a* (big squares) and *b* (big circles) between which the alleles can transition (small circles). Because alleles can reside in either of the two classes, allele *i* (doubled small circle at time *t*) is favoured by selection during [*t, t* + 1] if its class-specific fitness exceeds that of an average individual within its class. The change in frequency during this demographic time period due to selection is thus 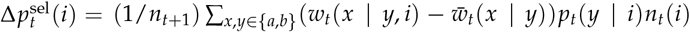 as given in eq. (4b). To obtain the frequency 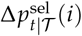 we must weight each offspring *i* produced due to selection during [*t, t* + 1] by their class-specific reproductive values giving the number of their descendants at *t*f under the neutralized process during [*t* + 1, *t*f], and then compare this to the total number of individuals at *t*f. This yields 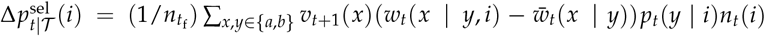 as given in eq. (8). **Panel B)** Evolutionary dynamics in a population without class structure where *wt*(*a* | *b, i*) = *wt*(*i*) for all *a, b* (Appendix A1.2.3). Allele *i* (doubled small circle at time *t*) is favoured by selection during [*t, t* + 1] if its fitness exceeds that of the average individual. The change in frequency during this demographic time period is 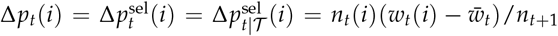. This can either be interpreted as the frequency of selected offspring produced during [*t, t* + 1] and measured at *t* + 1, or as the fraction of the total frequency of *i* individuals at the final time *t*f that descend under the average population process from all offspring produced due to selection during [*t, t* + 1] (see eq. A30 of the Appendix). These are equal because the average population process does not alter allele frequencies.

#### Disentangling Selection from Class Transmission

How then should one disentangle the effect of natural selection from the effect of class transmission when there are heritable differences in survival and reproduction in class structured populations, i.e. when *w*_*t*_(*a* | *b, i*) *≠ w*_*t*_(*a* | *b, j*) for different alleles *i* and *j*? We base the answer on two premises. First, because alleles can be found in different classes, natural selection on allele *i* should reflect heritable differential reproductive success by taking into account that each class can potentially make a distinct relative contribution to overall reproductive success (see Appendix A1.1 for how this premise is operationalized). Second, by definition of being an evolutionary force, natural selection should conserve allele frequencies so that a change in frequency due to selection of any one allele must be balanced by the change in other alleles segregating in the population. This is the conceptualisation of natural selection in the literature for populations without class structure, but in the presence of other forces acting on allele-frequency change (e.g., Nagylaki, 1992, eq. 2.7, p. 10-11, Frank, 1997, eq. 3, Grafen, 2000, Bürger, 2000, eq. 2.7, p. 125). We are then led to the following partitioning of allele-frequency change

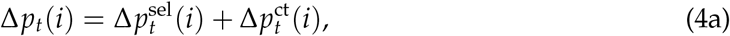

where the effect of selection on allele-frequency change is

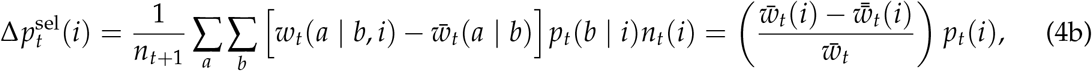

while the effect of class transmission is

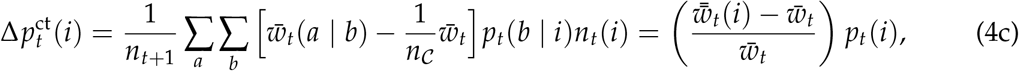

where 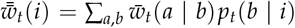 is the fitness of an allele *i* carrier if it were to reproduce as an average individual in each class, and recall that *n*_*C*_ is the number of classes in the population (see Appendix A1.1 for the derivation). We emphasize that this partitioning follows from the two premises stated above, hence different assumptions could lead to a different partitioning.

The first equality in eq. (4b) says that the effect of selection in each class is proportional to the difference between the class-specific fitness *w*_*t*_(*a* | *b, i*) of an individual carrying *i* as compared to the class-specific average fitness 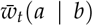 in that class, where the average in 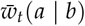 is calculated over all segregating alleles in the population. The class-specific average fitness 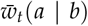 ensures that allele frequency is conserved under selection 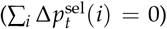, and can be considered as a reference fitness below and above which selection occurs in a class. Summing over all classes then produces the second equality in eq. (4b), which says that allele *i* is favoured by selection whenever it has a greater fitness than if it were to reproduce as an average individual within each class, i.e. 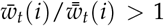. By contrast, the first equality in eq. (4c) says that the effect of class transmission in each class is proportional to the difference between the class-specific average fitness 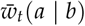 and the class-specific fitness in the absence of class-specific differences in reproduction 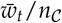 (where reproduction within and between classes is identical). Summing over all classes, allele *i* is thus favoured by class transmission whenever its fitness is greater if it were to reproduce as an average individual within a class as compared to an average individual in the total population, i.e. 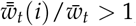, and this occurs whenever alleles are non-uniformly distributed across classes (see eq. A10 in Appendix A1.1). We emphasise that 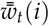 depends on the frequency *p*_*t*_(*b* | *i*) that is calculated under the full evolutionary process where *w*_*t*_(*a* | *b, i*) *≠ w*_*t*_(*a* | *b, j*). Whenever the process is neutral, i.e. *w*_*t*_(*a* | *b, i*) = *w*_*t*_(*a* | *b*) for all *i*, then 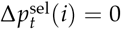 and 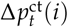 given by eq. (4c) reduces to eq. (3). A partitioning analogous to eq. (4) has been reached for a continuous time process in terms of phenotypic change [Lion, 2018a, eq. 2] as well as allele-frequency change [Priklopil and Lehmann, 2021]. Neither work however justified the partitioning based on the above-stated premises.

### Multigenerational Process in Class-Structured Populations

#### Framing Allele-Frequency Change Decomposition

Suppose now the process (eq. 1) runs over multiple generations between some initial time *t*_0_ and final time *t*_f_ where *𝒯* = [*t*_0_, *t*_f_] will denote this interval and contains the generation *t* discussed in eqs. (2-4). Our next aim is to ascertain how much of the total allele-frequency change 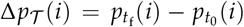 that has occurred during *𝒯* is due to natural selection and class transmission, and what fractions of this should be attributed to each generation of the multigenerational process. That is, we seek to find a decomposition

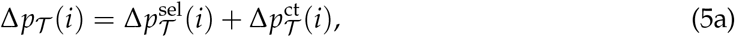

where

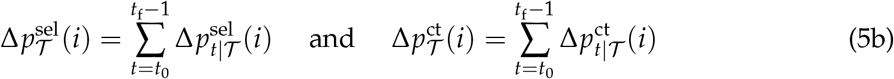

are, respectively, the cumulative fractions of allele-frequency change over *𝒯* that are attributed to selection and class transmission, and where 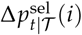 and 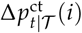 represent, respectively, the fractions of these changes that can be attributed to selection and class transmission occurring in generation *t*. The fraction of the total cumulative allele frequency change attributed to generation *t* is thus 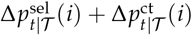.

Because 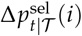 and 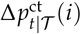 are assessed at the final time *t*_f_ of observation of the process and conditioned on the process starting at the initial time *t*_0_, they are not likely to match the partitioning in eq. (4) of the single-generational process that is conditioned on *t* and assessed at *t* + 1. Indeed, the intuition gained from eq. (4) already suggests that 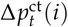 should not in general be equal to 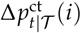 because 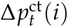 in eq. (4c) is conditioned on the state of the population at *t* and hence counts also offspring that are produced by parents who themselves were produced by selection in the previous generation(s) [*t*_0_, *t*]. Likewise, 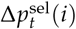 should not in general be equal to 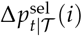 because the offspring produced in generation *t* due to selec-tion 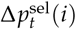 are produced into different classes, each of which may affect differentially their non-heritable reproductive success during [*t* + 1, *t*_f_] and hence may contribute differentially to 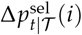. As a consequence, the fraction 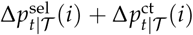 of the cumulative change attributed to generation *t* is also not in general equal to 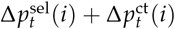. In the forthcoming section “Disentangling Selection from Class Transmission” we will make these arguments more precise. And so, our question is, how does one calculate the contributions in eq. (5)? To that end, we must first introduce the concept of an individual reproductive value.

#### Individual Reproductive Value

We now define 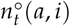 as the number of *a, i* individuals at time *t* that satisfy the recursion

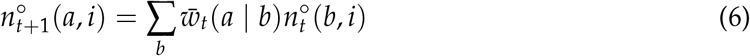

from *t*_0_ onward, with initial condition 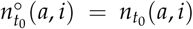 for all *a* and *i*. This recursion describes the change in the number of alleles in each class as if each allele in a given class were to reproduce and transition between classes as does an average individual from the same class, and whose fitness is 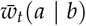. Because this class-specific average fitness 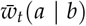 is calculated under the full observed evolutionary process with selection (i.e. the process defined by eq. 1), we refrain from calling the process defined by eq. (6) the ‘neutral process’ since this usually refers to a process where all individuals are exchangeable within a class [Ewens, 2004]. Instead, we call the process defined by eq. (6) the neutralized process, because all *i* follow eq. (6) and hence it is *as if* individuals are exchangeable under this process. The introduction of eq. (6) is motivated by the partitioning given in eq. (4) where 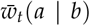 was shown to be the reference fitness that disentangles selection from class transmission. All variables with a superscript ◦ will then stand for variables that are determined by this neutralized process from time *t*_0_ onwards (Appendix A1.2). Formally, eq. (6) is the adjoint equation to the backward-in-time process

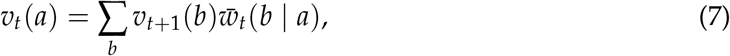

and we set the final condition to 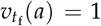 for all *a* (Box A). We can then interpret *v*_*t*_(*a*) as the *number of individuals alive at the final time t*_*f*_ *that under the neutralized process between* [*t, t*_*f*_], *where individuals survive and reproduce as average individuals within each class, descend from a single individual of class a alive at time t*. We call *v*_*t*_(*a*) the reproductive value of an (average) individual of class *a* (i.e. of a randomly sampled allele in class *a*). This is a representation of the classic notion of individual reproductive value in a finite-time process [Fisher, 1930, Grafen, 2006, Leslie, 1948, Taylor, 1990, Tuljapurkar, 1989, such a non-asymptotic reproductive value has also been called a ‘relative contribution’, Barton and Etheridge, 2011], which in a multi-type population process has indeed been evaluated from the average fitness within a class [Lessard and Soares, 2016, Lion, 2018a]. We denote by 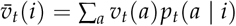 the average reproductive value of a carrier of allele *i*. We also define a population-wide average reproductive value 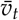 from 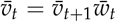 which is an adjoint equation to the population process 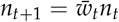 that we henceforth call the population-wide average (population) process. The interpretation of 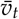 is the number of individuals at *t*_f_ that descend under the population-wide average population process from an average individual in the population at time *t* (i.e. 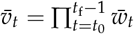).

#### Disentangling Selection from Class Transmission

We can now represent the contribution of selection attributed to generation *t* as

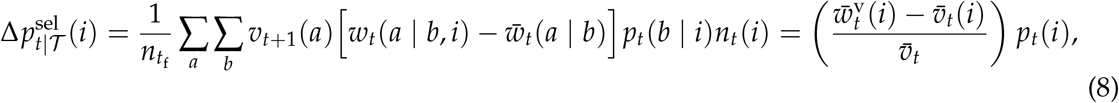

where 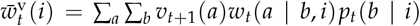 is the reproductive-value weighted average individual fitness giving the number of individuals at *t*_f_ that descend under the neutralized processes (eqs. 6-7) from all the offspring produced by a single individual *i* at *t* (Appendix A1.2). Because 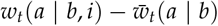 gives the additional number of offspring produced by *i* as compared to the average individual in the class during [*t, t* + 1], each of which leave *v*_*t*+1_(*a*) descendants under the neutralized process (eqs. 6-7), the summation over classes in eq. (8) isolates the one-generational effect of selection in an otherwise neutralized process. Note that by contrast to the change Δ*p*_*t*_(*i*) (eq. 4b) that assesses the offspring at time *t* + 1, the change 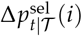 (eq. 8) assesses offspring at the final time *t*_f_ (see also Figure 2A).

**Figure 2:**
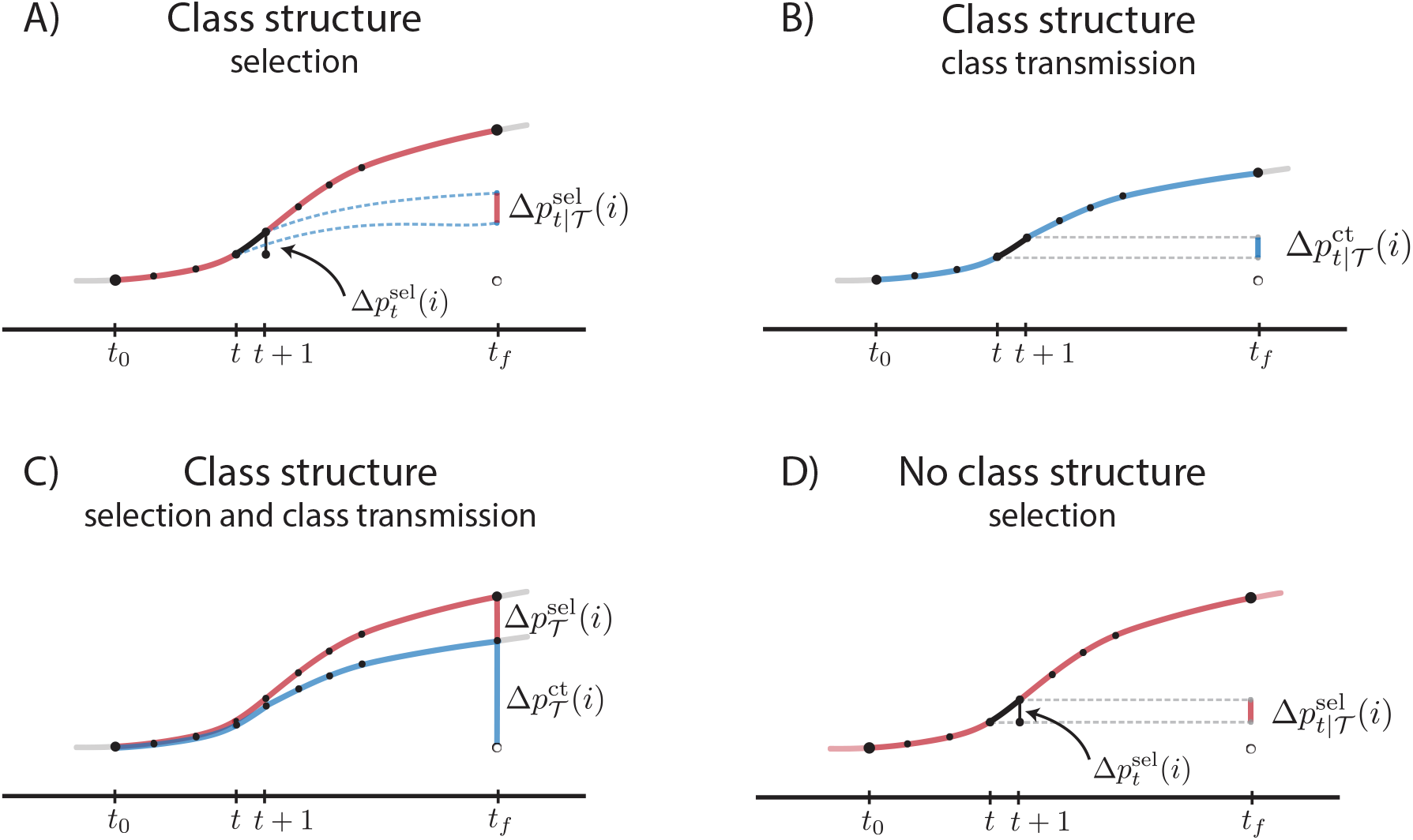
Partitioning of the allele-frequency change 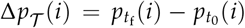 into fractions caused by selection 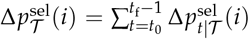 and class transmission 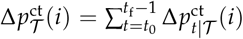. **Panel A)** Allele-frequency change due to selection 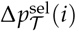 in a class-structured population. The dots along the thick curve indicate the observed allele frequencies in different genera-tions. The fraction 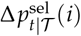 measured at time *t*f gives the difference between two frequencies (indicated by the short vertical line at time *t*f): the frequency of all *i* that underwent the full evolutionary process with selection until *t* + 1 and thereafter reproduced and survived under the neutralized process (top dashed line), and the frequency of all *i* that underwent the full evolutionary process only until time *t* and thereafter reproduced and survived under the neutralized process (bottom dashed line). Because the neutralized process can alter al-lele frequencies (the dashed lines are not horizontal), we have in general 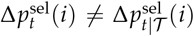 (compare eqs. 4b and 8). **Panel B)** Allele-frequency change Δ*pT* (*i*) due to class transmission in a class-structured population. The fraction 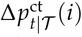 gives the difference between two fre-quencies (indicated by the short vertical line at time *t*f): the frequency of all *i* that underwent class transmission until *t* + 1 and where the excess offspring thereafter reproduced and survived according to the population-wide average population process (top dashed line), and the frequency of all *i* that underwent class transmission only until *t* and where the excess offspring thereafter reproduced and survived according to the population-wide average pro-cess (bottom dashed line). Because the population-wide average process does not alter allele frequencies these dashed lines are horizontal, and because we are conditioning on a (hypothetical) lineage of alleles that have underwent class transmission only, the dots along the thick curve are not observed frequencies in the population and must be calculated. This also implies that in general 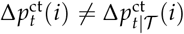 (compare eqs. 4c and 10), and explains why 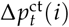 is not depicted. **Panel C)** The total multigenerational allele-frequency change Δ*p𝒯* (*i*) (top thick line) caused by selection 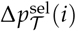 and class transmission 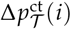 in a class-structured population. The sums 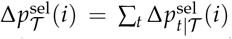 and 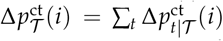 are obtained from panels B and C. **Panel D)** Allele-frequency change in a population without class structure where *wt*(*a b, i*) = *wt*(*i*) for all *a, b* (Appendix A1.2.3). The dashed lines are horizontal be-cause in populations without class structure we have 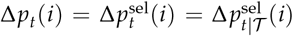, which is so because in the absence of selection during [*t* + 1, *t*f] the descendants reproduce according to the population-wide average population process and hence do not alter allele frequencies (Figure 1B; eq. A30).

In summary, eq. (8) is thus equal to the frequency of individuals measured at *t*_f_ that descend under the neutralized process from all offspring at *t* + 1 that were produced due to heritable differential reproduction and survival during [*t, t* + 1] by all parents *i* at *t*. This is the interpretation of the effect of selection attributed to generation *t* under in a multigenerational evolutionary process. Because 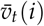 is always positive, selection favours allele *i* at *t* in a multigenerational process of span *𝒯* whenever

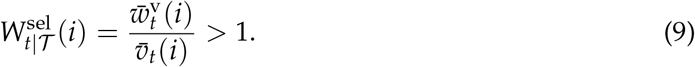

Here, 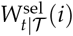 can be interpreted as the number of individuals at *t*_f_ that descend under the neutralized process from all offspring produced by a single *i* carrier at *t*, relative to the num-ber of individuals at time *t*_f_ that would descend from this individual under the neutralized process only. The fitness 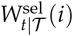 is thus the relative fitness attributed to selection at *t* in a multigenerational context.

Next, the contribution of class transmission that is attributed to generation *t* is

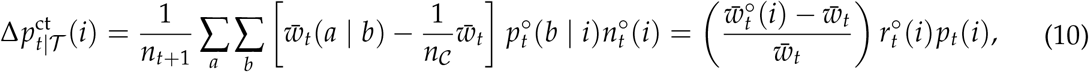

where 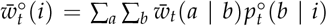 is the average fitness of an individual carrying *i* if it were to reproduce as an average individual of the class in which it resides at time *t* under the neutralized process (i.e. under eqs. 6-7). Such an individual itself descends from a lineage undergoing the neutralized process during [*t*_0_, *t*] since 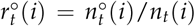 is the fraction of individuals *i* at *t* among those carrying *i* that descend from a lineage that has undergone only the neutralized process between [*t*_0_, *t*] (Appendix A1.2). Indeed, 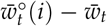 gives the average number of offspring at *t* + 1 produced by an individual *i* at *t* if it were to reproduce as an average individual in its class as compared to a population-wide average individual (compare with eq. 4c), which, when multiplied with the number 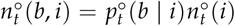 and divided by the total population size, produces 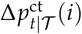 as required by the partitioning of total allele-frequency change (eq. 5). Note that in eq. (4c) we do not weight the offspring at *t* + 1 with reproductive values because we would then also count events of class transmission at later generations (and not only at the generation *t* of interest), and the parents contributing to class transmission at time *t* descend from individuals at the initial time *t*_0_ through class transmission only (see also Figure 2B).

In summary, eq. (10) is thus equal to the frequency of individuals measured at *t* + 1 that were produced due to class transmission during [*t, t* + 1] by all parents *i* that reach *t* through the neutralized process. This is the interpretation of the effect of class transmission attributed to generation *t* in a multigenerational evolutionary process. Because 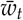 is always positive, class transmission favours allele *i* at *t* in a multigenerational process of span *𝒯* whenever

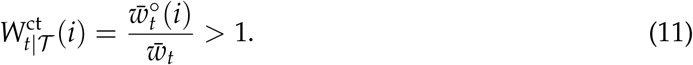

Here, 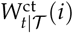 can be interpreted as the average number of offspring produced at *t* by a single *i* carrier if it were to reproduce as an average individual in each class and its ancestral lineage underwent the neutralized process, relative to the average number of offspring produced if this individual were to reproduce as an average individual in the population. The fitness 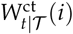 is thus the relative fitness attributed to class transmission at *t* in a multigenerational context.

#### Connection to Previous Work

We have produced a decomposition for allele-frequency change in a multigenerational process (eq. 5), where such change can be partitioned into generation-specific contributions caused by natural selection 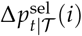 (eq. 8) and class transmission 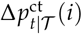 (eq. 10). These calculations are consistent with the partitioning of the single-generation effects of selection and class transmission, since eq. (4) is recovered by setting *t* = *t*_0_ and *t* + 1 = *t*_f_ in eq. (5) and eqs. (8)-(10), and they connect to a number of previous studies. To discuss these, let us denote by 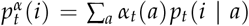 the class reproductive-value weighted allele *i* frequency at *t* where ∑_*a*_ *α*_*t*_(*a*) = 1 (see eq. A19 for the representation of *α*_*t*_(*a*)). Then, the fraction of the total cumulative allele-frequency change attributed to selection in generation *t* is exactly the class reproductive-value weighted change at *t*:

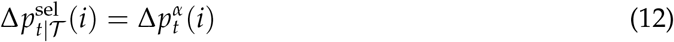

(eq. A19). The right-hand side of eq. (8) is therefore conceptually equivalent to eq. (10) of Grafen [2015] for allele-frequency change in a panmictic population, to eq. (7) of Lion [2018a] for the phenotypic change in a panmictic population, to eq. (5) of Lion and Gandon [2022] for the allele-frequency change in periodically fluctuating environments, and to eq. (69) of Priklopil and Lehmann [2021] for allele-frequency change in a spatially structured population. This previous work, however, did not provide the biological interpretation of the change 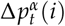 in terms of contributions to final frequencies in a (finite or infinite-time) multigenerational process, i.e. it did not discuss the left-hand side of eq. (12), which is interpreted as the change in the arithmetic average allele frequency due to selection in generation *t* but assessed at the end of the observed evolutionary process. Also, it did not provide the corresponding expression for the allele-frequency change 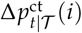 caused by class transmission. The cumulative contribution of selection for phenotypic change (here corresponding to 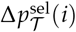) was in turn discussed in Gardner [2015]. However, this previous work focused on the large-time behavior of populations (*t*_f_ → ∞), and moreover, it did not define operationally the reproductive-value weights and instead appealed to Taylor [1990] who considers a reference fitness that is subject to a neutral process only (i.e. it is the fitness in a monomorphic population for the wild-type or resident allele). The recursion for reproductive value eq. (7) should however be expressed in terms of the average class-specific fitness (the fitness of a randomly sampled allele in each class) in the observed focal population subject to selection as explicitly emphasized in Lessard and Soares [2016], Lion [2018a], and so it is unclear to what type of evolutionary process the formalization of Gardner [2015] really applies to. Likewise, the cumulative change 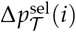 was considered in Lehmann and Rousset [2014, footnote 3], but here too, the reproductive-value weights were not operationalised. We here ascertained that the neutralized process must be evaluated under the (within-class) average population process (eq. 6) for 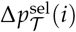 to represent the cumulative contribution of selection in the observed population.

In a series of two-allele models under weak selection induced by weak phenotypic deviations, the cumulative contribution of selection was considered in populations of small (finite) size under the form of the probability of fixation [Leturque and Rousset, 2002, Rousset and Ronce, 2004]. Under weak selection, it is sufficient to evaluate reproductive values under the neutral process determined by the wild-type or resident population only (see Rousset, 2004 for applications to general demographic situations). Furthermore, and finally, in large populations and under weak phenotypic deviation, the effect of class transmission vanishes asymptotically (as *t*_f_ → ∞). In this case for sufficiently large *t* the reproductive-value weighted allele-frequency change is approximately equal to the change in the arithmetic allele frequency [Priklopil and Lehmann, 2021], namely

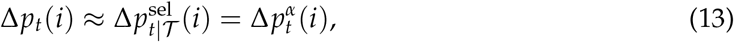

which thus fully characterizes allele-frequency change in a class-structured population (see also Appendix A3.2). Here too, it is sufficient to evaluate the reproductive values under the neutral process determined by the resident population only (e.g., Appendix A3.2). The term 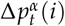 has long been in use in phenotypic models to describe reproductive-value weighted allele-frequency change under weak selection [e.g., Lion, 2018a, Priklopil and Lehmann, 2021, Rousset, 2004, Roze and Rousset, 2004, Stubblefield and Seger, 1990, Taylor, 1990, 1996]. Eq. (13) thus connects this allele-frequency change under weak selection back to the final frequencies and the full frequency change at generation *t* of a multigenerational process. An expression analogous to 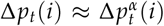 was also reached in terms of phenotypic change in panmictic populations [Barfield et al., 2011, Lion, 2018a,b]. Finally, the expression 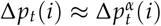 holds also for evolution in spatially subdivided class-structured populations [Priklopil and Lehmann, 2021].

### Geometric Mean Fitness

Having identified the contributions of selection and class transmission in a multigenerational process, it seems natural to ask whether one can summarize the evolutionary process by a single number – a representation of mean fitness – that would predict whether the allele frequency increases or decreases as a result of natural selection and/or class transmission? In Appendix A2 we show that evolution favors an increase of allele *i* over the entire time-interval *𝒯* whenever 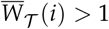 with

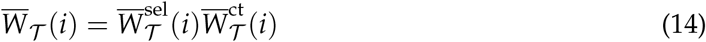

being the geometric mean fitness of allele *i* where

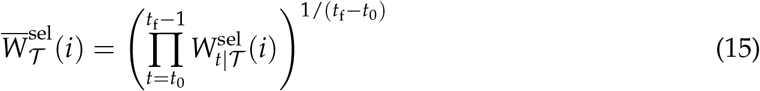

and

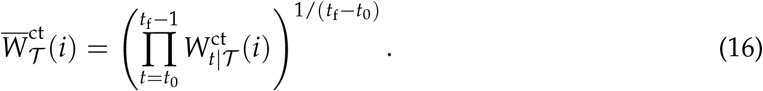

where 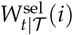 and 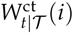 are as in eqs. (9) and (11), respectively. As discussed above (section Disentangling Selection from Class Transmission), selection favours allele *i* in generation *t* whenever 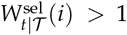, and class transmission favours allele *i* in generation *t* whenever 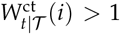 (see also Appendix A2). Over the entire time-interval *𝒯*, selection thus favours allele *i* whenever 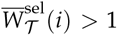, while class transmission favours allele *i* whenever 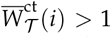. We have thus obtained a partitioning of the geometric mean fitness into the means 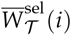 and 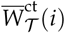 summarizing the multigenerational contribution of selection and class transmission for allele-frequency change, respectively, both of which are grand means since the averages therein involve averaging (arithmetically) expected number of offspring over all classes within a generation and (geometrically) over all generations within the time-frame of interest.

Importantly, it is not a conclusion that if allele *i* is favored by evolution, 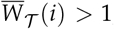, it is also favored by selection, 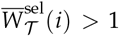, nor vice versa. This suggests that one should worry about the effect of class transmission on the evolutionary process if one is interested in un-derstanding the exact effect of selection in class-structured populations. Yet the theoretical literature routinely uses representations of mean geometric fitness to characterize evolutionary stable population states thus attributing such effects entirely to selection (e.g., Caswell, 2001, Ferrière and Gatto, 1995, Svardal et al., 2015). These representations are justified because they consider the asymptotic spread of a rare allele in the population (Box B; Appendix A3.1). In this case, selection is the determining evolutionary force because class structure converges in a large time limit to its steady state and will no longer contribute to allele-frequency change. The same is true for the characterization of the dynamics of alleles that express closely similar phenotypes leading to weak selection (recall eq. 13, see also Appendix A3.2). Finally we note that one can also express 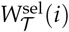 in terms of the 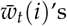, but then the generation-specific partitioning is incorrect and one must nevertheless define it relative to the same reference fitness as in eqs. (14)-(16) to account for the effect of class transmission (Appendix A2.2).

## Discussion

We analyzed allele-frequency change in a class-structured population with the aim to decompose the changes caused by natural selection and the process of class transmission allowing for frequency-dependence, density-dependence, and time-dependent environmental changes. Three main messages arise from our analysis.

First, we presented the standard representation for the single-generation change in allele frequency (eq. 2). We then calculated the contributions of selection and class transmission to this change (eq. 4) and showed that selection acting on an allele must be defined relative to the average fitness in each class (eq. 4b). These single-generation contributions of selection and class transmission, as our analysis reveals, should not however be used as net contributions to allele-frequency change in a multigenerational context where the measurements are taken over multiple time steps (Figures 1 and 2). This is so because, on one hand, the contribution of class transmission contains the contributions of selection in the past, and hence in a multigenerational context it conflates the effect of class transmission with past effects of selection. On the other hand, the contribution of selection fails to take into account that the descendants of the selected offspring undergo class transmission in the future thus altering the magnitude of this allele-frequency change, and hence does not take into account that non-heritable survival and reproduction differences affect the evolutionary process in class-structured populations. Hence, the single-generational allele-frequency change that is still widely used to describe the essence of natural selection (e.g., the review of Queller, 2017 p. 347) does not generally capture the net contribution of selection in a multigenerational context in class-structured populations. Yet all allele frequency change leading to adaptation is certainly multigenerational and populations are generally class structured.

Second, we provided the expressions for the contribution of selection and class transmission attributed to generation [*t, t* + 1] in a multigenerational process spanning between some initial time *t*_0_ and final time *t*_f_ (eqs. 8–10). We showed that these contributions must be de-fined in the context of the entire span of the evolutionary process under focus and requires a careful account of the notion of reproductive value for which we provided a biological definition in the context of a finite-time process (see eq. 7). Based on our analysis we suggest the following definition for the effect of selection acting on allele *i* and attributed to a specific demographic time step [*t, t* + 1] of interest: it is the difference between two frequencies measured at the end of the evolutionary process at time *t*_f_, (1) a frequency of allele *i* if it were to descend from a lineage in a population that until *t* + 1 first underwent the full evolutionary process with selection and class transmission, but thereafter selection was ‘turned off’ and the population underwent only class transmission (induced by the neutralized process), and (2) a frequency of allele *i* if it were to descend from a lineage in a population that underwent the full process only until *t* after which the population underwent class transmission (Figure 2). This definition thus isolates the one generational effect of selection during [*t, t* + 1] from an otherwise neutralized (or non-selective) process of class transmission. Because these effects attributed to generation [*t, t* + 1] are assessed at *t*_f_ they are in general different to evolutionary changes in the single-generation context assessed at *t* + 1 (eq. 4b). It may therefore be the case that while the single-generation effect of selection contributes negatively to allele-frequency change, the multigenerational effect may be positive and vice versa. For weak selection resulting from individuals expressing closely similar phenotypes, a simpler picture emerges as class-transmission becomes negligible in distant future. Then, all allele-frequency change in the present is captured by the reproductive-value weighted allele-frequency change, and this also gives the present effect of selection in terms of a frequency assessed in the distant future. In this case the reproductive-value weighted allele-frequency can be interpreted as a measure that aligns the current effect of selection with both current and future allele-frequency change thereby straddling single-generational and multigenerational perspectives of selection (eq. 13).

Third, we summarized the contributions of selection and class transmission over the entire evolutionary process in terms of geometric mean fitness’s (eqs. 15-16). The representation of the (geometric) mean fitness of a particular allele summarizing the multigenerational effect of selection is the relative reproductive-value weighted average individual fitness of a carrier of that allele (eq. 15). This gives the grand mean – a mean taken over all classes within a generation and over all generations of the evolutionary process under interest – of the expected number of individuals that descend under the neutralized process from the expected offspring number produced by a single carrier of the allele under focus, relative to the grand mean of the expected number of individuals that descend from that carrier under the neutralized process only. While this fitness is generally not sufficient to predict the direction of evolution, it should do so over long timescales (*t*_f_ → ∞) where the class-transmission often vanishes. In particular, this is the case asymptotically and underlies the widespread use of invasion fitness as a representation of fitness summarizing the multigenerational effect of selection (see Box B). This is also the case in populations where class-transmission is a much faster process than selection, as for instance under weak selection. In both cases the class-specific genetic structure converges fast under class transmission to its steady state and the long-term dynamics is governed purely by selection (see Appendix A3.1 for details). In biological scenarios where these two assumptions are not met exactly, we also expect that the geometric mean for selection calculated over sufficiently many generations gives a good approximation for the effect of selection and can be used to predict the evolutionary trajectory of an allele beyond the time-frame of observation.

We formulated all these results for haploid asexual populations, but our analysis can in a straightforward manner be generalized to more complex life-cycles and sexually reproducing populations. Likewise, we considered only panmictic populations, but the main results are bound to carry over to spatially structured populations. Our interpretations of allele frequency changes due to natural selection in class-structured populations should thus apply regardless of the details of the underlying population processes. Overall, we hope to have given a satisfactory answer to the question posed in the introduction and clarified the operation of natural selection in class-structured populations.

## Acknowledgments

We thank the the four reviewers for their useful comments, as well as the Associate Editor Sébastien Lion and the Editor Erol Akçay for their constructive suggestions that markedly improved the presentation of this paper.

## A1 Allele-Frequency Change in Class-Structured Populations

### A1.1 Single Generational Process

We here derive eqs. (2)–(4). First, note that

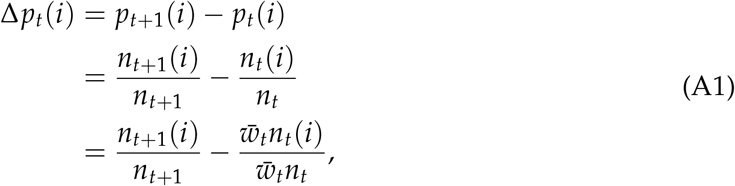

where the second equality follows from the definition of allele frequency. By substituting 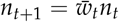 (obtained by summing eq. 1 over all classes and alleles) we obtain

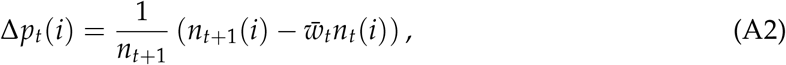

and by substituting 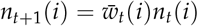 (obtained by summing eq. 1 over all classes) yields

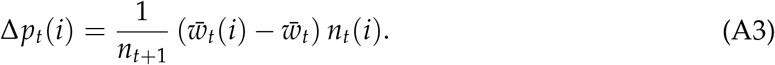

This is the first equality in eq. (2) and the second therein is obtained by substituting for 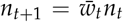 in the denominator. Using 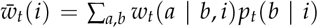 then allows to write eq. (A3) as

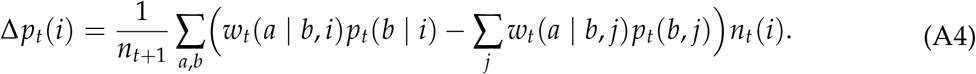

Now suppose that all alleles have exactly the same fitness, i.e. *w*_*t*_(*a* | *b, i*) = *w*_*t*_(*a* | *b*) for all *i*, then this allele-frequency change reduces to

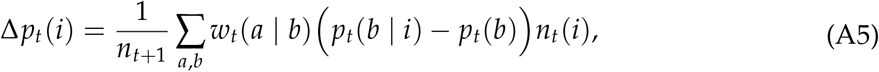

where we used *p*_*t*_(*b*) = ∑_*j*_ *p*_*t*_(*b, j*) and which shows eq. (3).

Next, we consider the case where *w*_*t*_(*a* | *b, i*) *≠ w*_*t*_(*a* | *b, j*) and show that selection acting on an individual carrying allele *i* in a given class must be defined relative to the average individual fitness within that class. To do this, we here expand on the two premises given in the main text to conceptualise natural selection. First, natural selection is an evolutionary force changing the relative numbers of alleles, i.e. their frequencies, and this is as a result of differential reproductive success caused by heritable genetic differences. In class-structured populations, however, alleles may reproduce and/or survive differentially also due to non-heritable class-specific differences, but because non-heritable differences do not contribute to the selective process these should be disregarded. This is motivated and consistent with previous literature about evolution in class-structured populations ([e.g., Grafen, 2006, Leturque and Rousset, 2002, Lion, 2018a, Stubblefield and Seger, 1990, Taylor, 1990]) and overall entails that selection should reflect that each class can potentially make a distinct relative contribution to overall reproductive success due to heritable differences. This leads us to operationalize our first premise by letting the average excess of *i* individuals in class *a* produced by *i* carriers in class *b* due to selection be proportional to *w*_*t*_(*a* | *b, i*) − *f*_*t*_(*a* | *b*), where *f*_*t*_(*a* | *b*) is a reference measure of reproductive success to be derived from the analysis. This reference measure is introduced precisely to tease out from the selective process any differences in reproduction that are due to non-heritable class-specific differences, and as this measure must be the same for all alleles we do not index it by *i*. Second, natural selection is a conservative evolutionary force, i.e. it maintains total allele frequency so that if allele *i* increases in frequency, 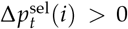, another must decrease as a result of natural selection and the total allele allele frequency must be zero sum: 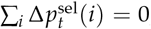, which is the standard conceptualisation of natural selection in the theoretical literature (e.g., Nagylaki, 1992, eq. 2.7, p. 10-11, Frank, 1997, eq. 3, Grafen, 2000, Bürger, 2000, eq. 2.7, p. 125).

With these premises in mind, let us partition the number 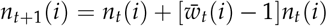 of alleles *i* produced trough survival and reproduction in the population as

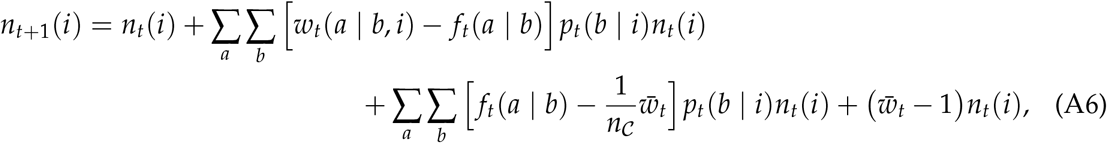

where the first summand gives the change in *i* number caused by differential reproductive success due to allele differences, the second gives the change in *i* number caused by individuals residing in different classes, and the last term gives a change due to the population-wide average process (population growth). This last term should not contribute to allele frequency change and indeed by inserting eq. (A8) into eq. (A2) produces

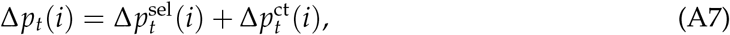

with

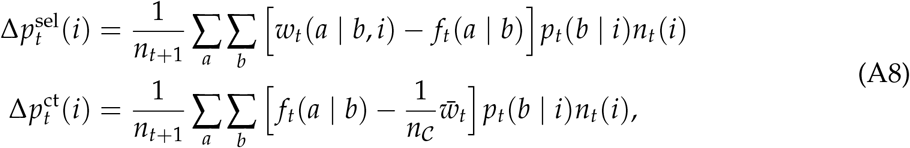

which give the changes in frequency of *i* caused by selection (‘sel’), with yet to be established *f*_*t*_(*a* | *b*), and class transmission (‘ct’). Next, summing over *i* in the first line of eq. (A8) produces

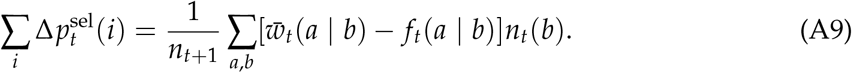

Because natural selection is conservative 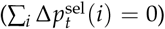, we must have 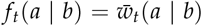 for all *a, b* due to the fact that class-specific differences may be independent, which implies 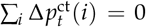 and therefore proves eq. (4). We finally note that by substituting 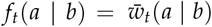 into the second equation in eq. (A8) and re-arranging we get

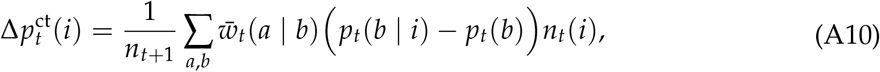

thus confirming eq. (A5), saying that class transmission is induced by the non-uniform distribution of alleles across classes.

When there is no class structure, i.e. *w*_*t*_(*a* | *b, i*) = *w*_*t*_(*i*) for all *a* and *b*, we have 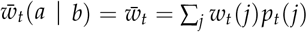. Then, 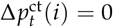 and eq. (A7) reduces to

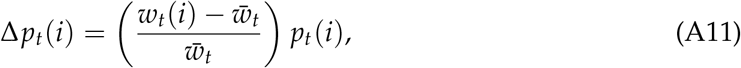

which is the standard representation of allele-frequency change in populations without class structure (Gillespie, 1991, eq. 4.1, Nagylaki, 1992, eq. 2.8).

### A1.2 Multigenerational Process

We here derive eq. (8) and eq. (10). To that end, we define *n*_*t*|*h*_(·) and *p*_*t*|*h*_(·) as the state variables at *t* that underwent the full evolutionary process between [*t*_0_, *h*] and that from time *h* onward have undergone the neutralized process where individuals reproduce as average individuals in each class. In the main text, we use the short-hand notation 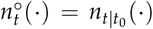 and 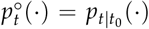 for variables that have undergone the neutralized process right from the onset (i.e. the initial time *t*_0_), but we must define the more general state variables in this appendix to derive the results. In particular, *n*_*t*|*h*_ (*a, i*) gives the number of individuals *a, i* at time *t* that descend from a lineage of individuals that underwent the full evolutionary process between [*t*_0_, *h*] and the neutralized process between [*h, t*]. For *h* ≤ *t* < *t*_f_, this state variable thus satisfies the recursion

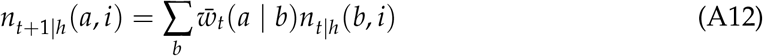

(compare with eq. 6 in the main text) with initial condition *n*_*h*|*h*_ (*a, i*) = *n*_*h*_(*a, i*), which is the final state of the state variable *n*_*t*_(*a, i*) that has undergone the full evolutionary process (eq. 1) for *t*_0_ ≤ *t* ≤ *h*. Next, we introduce the state variable

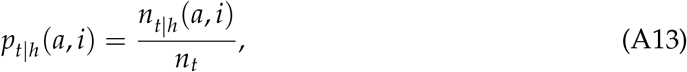

which gives the proportion of individuals in the population at time *t* that descend from a lineage of *a, i* individuals that underwent the full evolutionary process between [*t*_0_, *h*] and that from time *h* onward have undergone a neutralized process (and recall that *n*_*t*_ = *n*_*t*|*t*_ and hence this proportion is relative to the total number of individuals under the full evolutionary process). We can equivalently interpret *p*_*t*|*h*_ (*a, i*) as the frequency of individuals of type *a, i* at the final time *t*_f_ that descend from a lineage of individuals that underwent selection and class transmission between [*t*_0_, *h*] and then only class transmission between [*h, t*] (relative to the number of individuals that underwent the full evolutionary process from *t*_0_ onward).

#### A1.2.1 Disentangling Selection from the Evolutionary Process

We are now ready to derive eq. (8). First, by using eq. (A13) we observe that in a multigenerational process over *𝒯*, the effect of selection in generation *t* on allele frequency can be defined for all *t* as

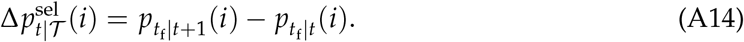

Indeed, the variable 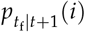 gives the frequency of individuals *i* at *t*_f_ that descend from individuals that underwent selection and class transmission until *t* + 1 and then class transmission until *t*_f_. Hence, subtracting from 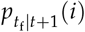, the allele frequency 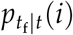 where selection acts only until time *t*, we isolate in eq. (A14) one iteration of selection [*t, t* + 1] from an otherwise class-transmission process during [*t, t*_f_]. Applying eq. (A14) recursively, we get

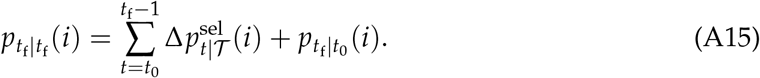

Then, by using the consistency relation 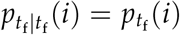 and by reorganizing the terms gives

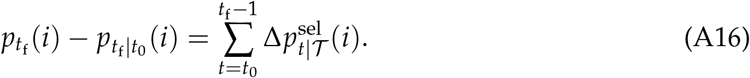

Because 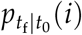 is the frequency of *i* at time *t*_f_ if only class transmission was in play over the entire *𝒯* (and 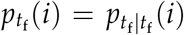 if both selection and class transmission was in play), we have obtained that 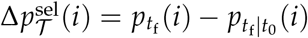 gives the fraction of the total allele-frequency change that is due to selection with 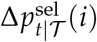 as the contribution of selection in generation *t*.

Next, we evaluate 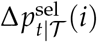 as defined by eq. (A14). For this, we note the identity

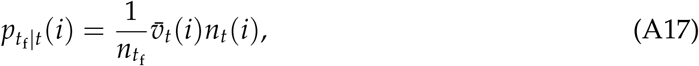

where we used the definition for reproductive values (eq. 7) and the average reproductive value

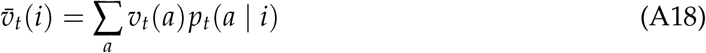

of an individual carrying allele *i* at time *t* as defined in the main text. Notice that eq. (A17) can also be written as

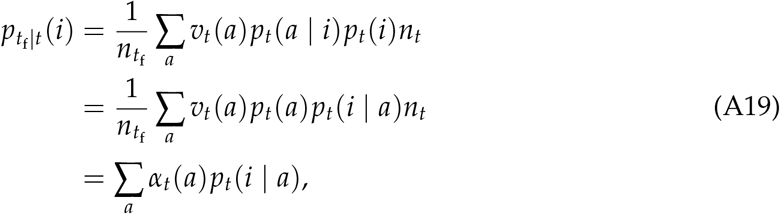

where 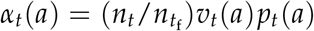 is the class reproductive value satisfying ∑_*a*_ *α*_*t*_(*a*) = 1 since 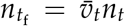. Hence, 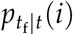 is the (class) reproductive-value weighted allele frequency used in the evolutionary analysis of class-structured populations (e.g., Lion, 2018a, Priklopil and Lehmann, 2021, Rousset, 2004, Rousset and Ronce, 2004, Taylor, 1990, 1996).

Now, the weighted allele frequency (eq. A17) satisfies

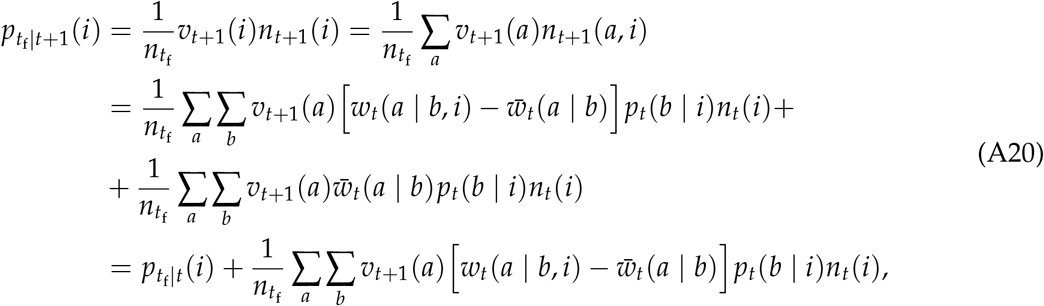

where we applied eq. (A18) in the second equality, eq. (1) in the third equality, and eq. (7) and eq. (A17) in the fourth equality. From the definition in eq. (A14) this thus proves the first equality in eq. (8).

The second equality in eq. (8) can be obtained by noting that 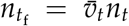 where 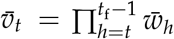, in which case we can write eq. (A17) equivalently as

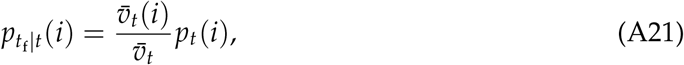

and hence

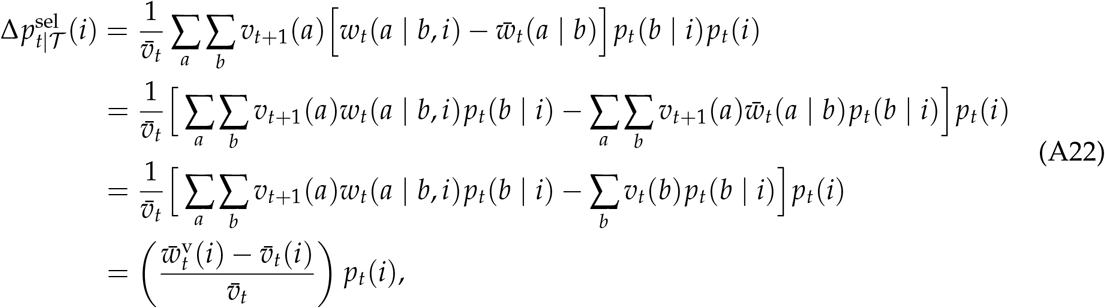

where in the third equality we applied eq. (7) and in the fourth equality the definition of 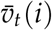

and

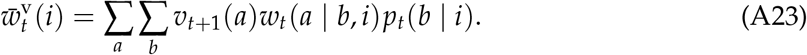

#### A1.2.2 Disentangling Class Transmission from the Evolutionary Process

We here derive eq. (10). We observe that in a multigenerational process over *𝒯* the effect of class transmission in generation *t* on allele frequency can be defined for all *t* as

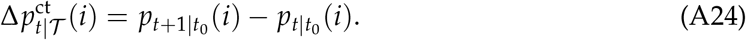

Indeed, 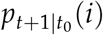 gives the frequency of *i* individuals that underwent class transmission until *t* + 1, and hence by subtracting from this the frequency 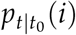 of *i* individuals that underwent class transmission until *t*, we isolate one iteration of class transmission during [*t, t* + 1]. Now, by setting *t* = *t*_f_ − 1 into eq. (A24), we have

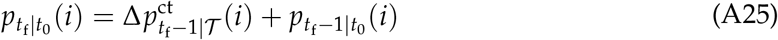

and a recursive substitution produces

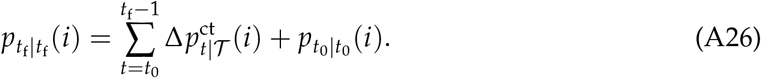

Then, by noting that 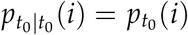 and by reorganizing the terms gives

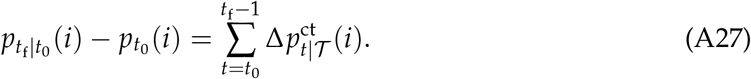

Because 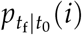 is the frequency of *i* at time *t*_f_ if only class transmission was in play over the entire time-interval *𝒯*, we have obtained that the total change in allele frequency due to class transmission is 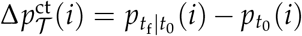 and 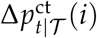 is the contribution of class transmission in generation *t*.

Next, we evaluate 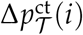 as defined by eq. (A24). Using eq. (A13), we can write the recursion

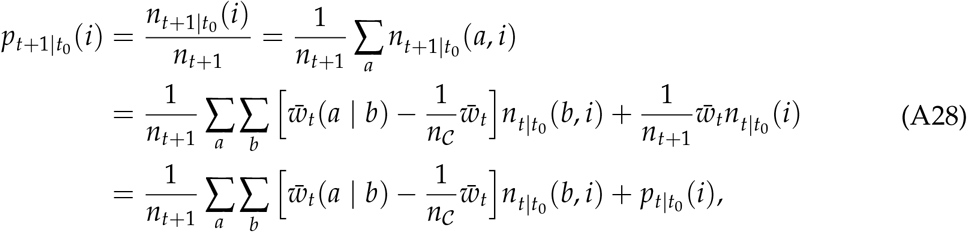

where in the third equality we used eq. (A12) (eq. 6 in the main text) and in the fourth equality 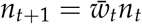 and eq. (A13). Reorganization and using 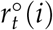 as defined in the main text, as well as recalling the shortcut notation 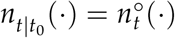 and 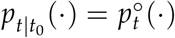 yields eq. (10).

#### A1.2.3 Reduction to Population Models Without Class Structure

We now show that our derivation is consistent with the standard population genetic model without class structure where *w*_*t*_(*i*) = *w*_*t*_(*a* | *b, i*) for all *a, b*. Substituting this into eq. (10) gives 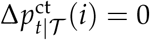 and when substituting it into eq. (8) yields

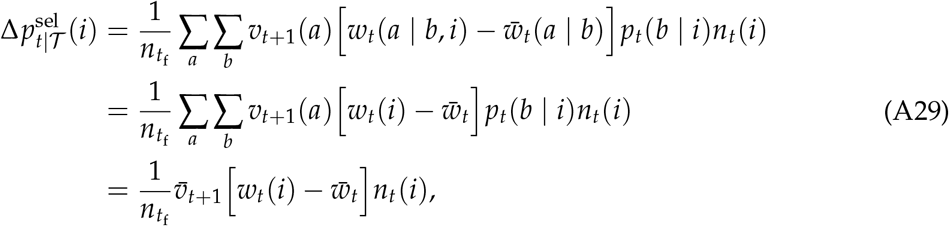

where we used 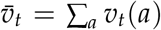 and 1 = ∑_*b*_ *p*_*t*_(*b* | *i*). Using 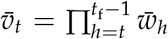 we can write 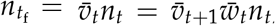, and substituting into eq. (A29) shows that the fraction of allele-frequency change attributed to selection in generation *t* is equal to the right-hand side of eq. (A11) and thus we have in force of eq. (A7) (but where class transmission has no effect) that for populations without class structure

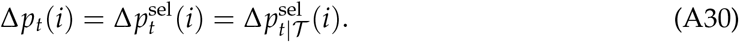

However, in a multigenerational context, we can interpret 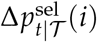 as the fraction of the total frequency of *i* measured at the final time *t*_f_ that is attributed to selection in generation *t*, where all descendants of the average excess number of individuals produced over [*t, t* + 1] undergo the population-wide average process in [*t* + 1, *t*_f_]. This is a novel interpretation of allele frequency change in the standard model and justifies the use of reproductive values in Figure 1B.

## A2 Geometric Mean Fitness

### A2.1 Geometric Mean in terms of multigenerational relative fitness

We here derive eqs. (9), (11) and eqs. (14)-(16). First, we calculate eqs. (9) and (15). Using eq. (A13) we obtain the recursion

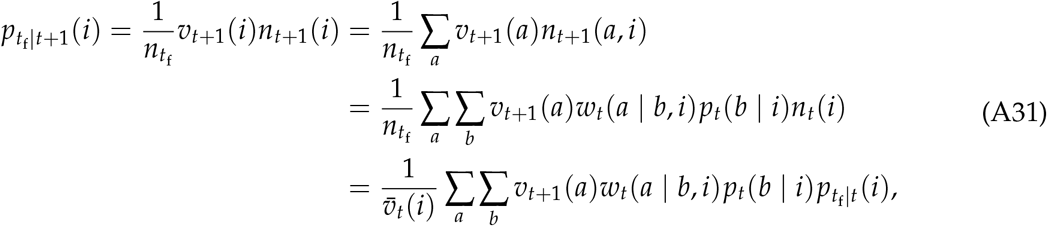

where in the third equality we used eq. (1) and in the final equality we used the relation 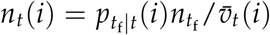, which was obtained from 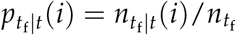 (using eq. A13) and the identity 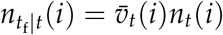. Then, using the definition of 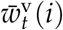 produces

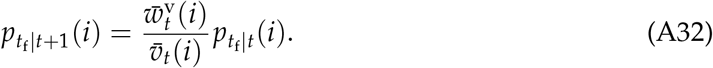

Because 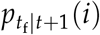 is the frequency of *i* at *t*_f_ that descend from the lineage of *i* in a population that underwent selection and class transmission until *t* + 1 and thereafter only class transmission, and 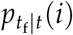 is the frequency of *i* at *t*_f_ if selection acted only until *t*, then 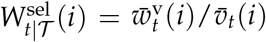 is the relative fitness attributed to only selection in generation *t*. Setting *t* + 1 = *t*_f_ on the left hand side in eq. (A32) and iterating backward in time yields

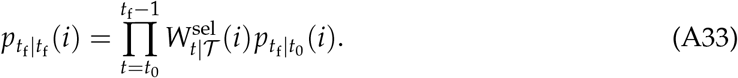

Because 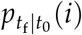 is the allele frequency at time *t*_f_ in a process without selection and 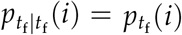 is the frequency at *t*_f_ with selection, the product in eq. (A33) summarizes the fitness excess over *𝒯* caused by selection only. Taking the geometric mean of the product in eq. (A33) then together with eq. (A32) proves eqs. (9) and (15).

Second, we calculate eqs. (11) and (16). Consider the recursion

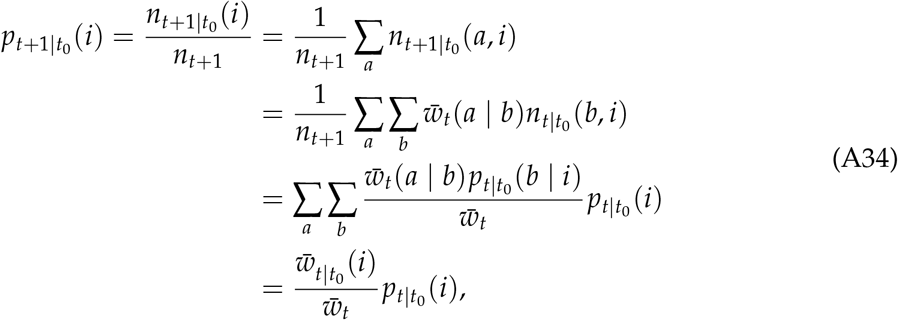

where we used eq. (A12) (eq. 6 in the main text) and

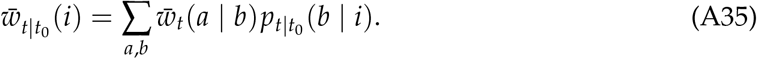

This is the fitness of a single individual *i* at *t* coming from a lineage of individuals that descend under the neutralized process and will be different for different alleles if alleles are initially distributed differently between classes: 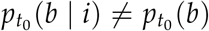. Note that in the main text we used the shorthand notation 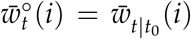. Because 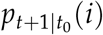 is the frequency of *i* at *t* + 1 that descend from a lineage of *i* that underwent class transmission between [*t*_0_, *t* + 1] and 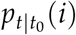 is the frequency of *i* at *t* whose lineage underwent class transmission between [*t*_0_, *t*], then 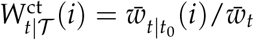 is the relative fitness attributed to only class transmission in generation *t*. Setting *t* + 1 = *t*_f_ on the left hand side in eq. (A34) and iterating backwards in time yields

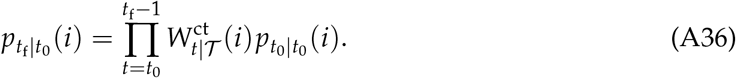

Because 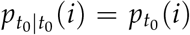 is the initial allele frequency at *t*_0_ and 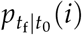 is the final frequency under class transmission only, the product in eq. (A36) summarizes the fitness excess over *𝒯* caused by class transmission (see below further discussion on the relative fitness attributed to class transmission in generation *t*). Taking the geometric mean of the product in eq. (A36) then together with eq. (A34) proves eqs. (11) and (16).

Finally, we calculate eq. (14). Substituting eq. (A36) into eq. (A33), and by using 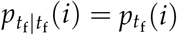 and 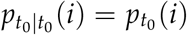, we get

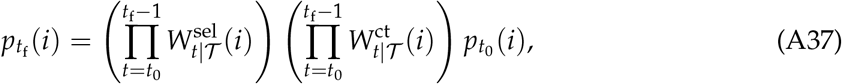

where 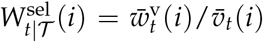 and 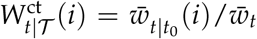. Taking the geometric means proves eq. (14).

As noted above, 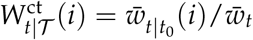 in eq. (A37) is the relative fitness for class transmission of a single *i* individual at *t* in a multigenerational process over *𝒯*, given we have sampled an individual at *t* who is itself a product of class transmission. If we want to condition on the realized population state at some later generation *h*, we can use the fact that

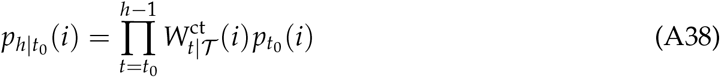

and substituting this into (A37) we have

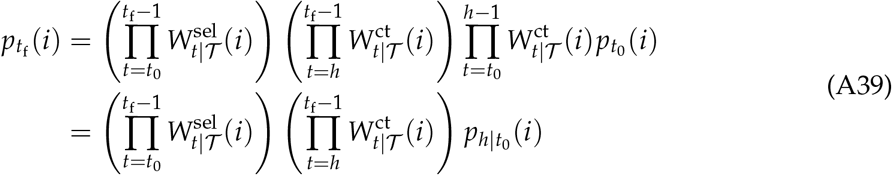

and using the identity 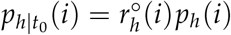 we get

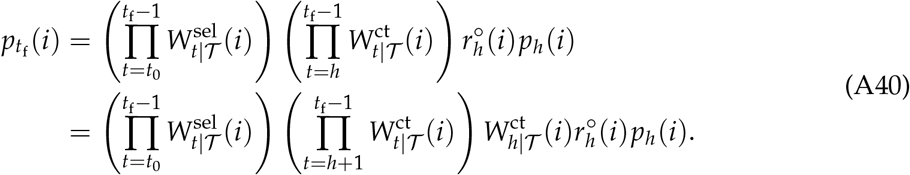

This shows that 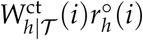 is the relative fitness for class transmission of a single randomly sampled *i* individual in generation *h*.

### A2.2 Alternative Representation of Geometric Mean for Selection

Here we show that one can write the geometric mean for selection (eq. 15) also in terms of individual fitness functions as

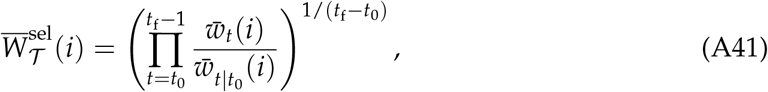

and recall that in the main text we use the shorthand notation 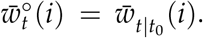. To show this, we first observe that the cumulative fitness of *i* over *𝒯* caused by selection only is 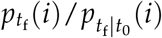 (compare with eq. A16). To obtain 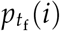, we can use the recursion 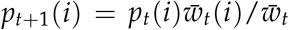 recursively over *𝒯* and we get

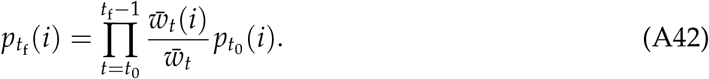

Now, substituting this and the frequency 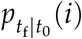 from eq. (A36) into 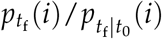, and taking the geometric mean, gets us eq. (A41).

Next, we show that eq. (A41) and eq. (15) are equivalent. To do this, we first derive a recursion for *p*_*h*|*t*_(*a* | *i*) over [*t, t*_f_] as

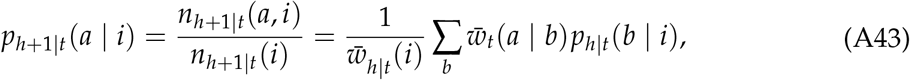

where we applied eq. (A12) (eq. 6 in the main text) and 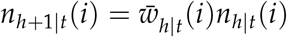 with 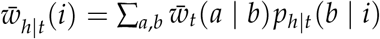, and where the initial condition satisfies *p*_*t*|*t*_(*a* | *i*) = *p*_*t*_(*a* | *i*). Then, we produce

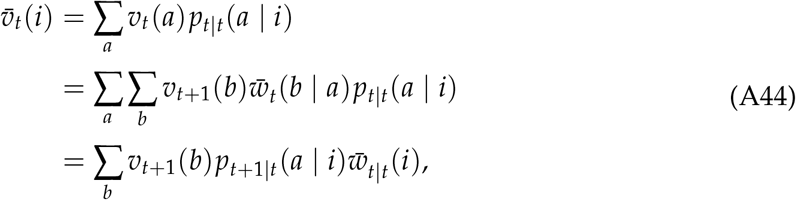

where the first equality follows from the definition of 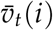, and in the second equality we used eq. (7) and in the third equality we used the relation 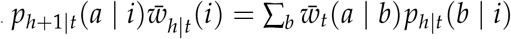 obtained from eq. (A43). Applying the logic in eq. (A44) recursively we get

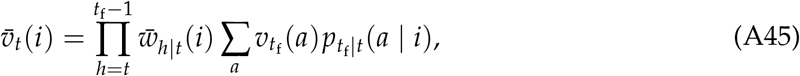

and by noting that 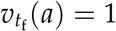 for all *a* and 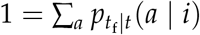 produces

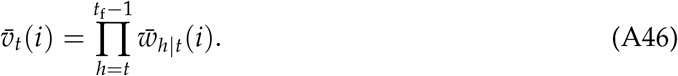

We can then write

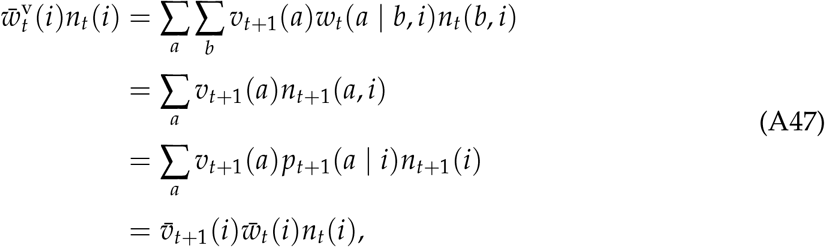

where the first equality follows from the definition of 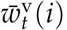, and in the second equality we used eq. (1) and in the final equality we applied the definition of 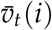 and 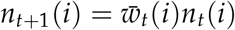. We have thus produced the identity

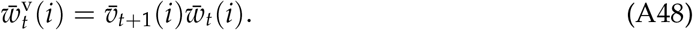

We can substitute this into

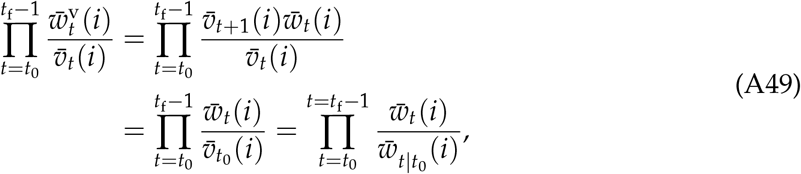

where in the second equality we used 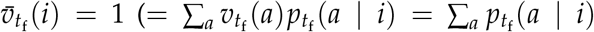 by assumption), and in the final equality we applied eq. (A46). This shows that the eqs. (15) and (A41) are indeed equivalent.

If one is only interested in the net geometric mean, then one can use eq. (A41) instead of eq. (15). In fact, eq. (A41) may in some situations be easier to apply because each factor 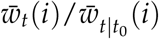 only depends on the past events whereas in eq. (15) each factor 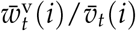 requires calculating also the future states of the population via reproductive values. However, the per generation contributions 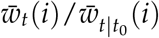 in eq. (A41) and 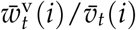 in eq. (15) are not equal, and only the latter is the correct fitness describing the effect of selection in generation *t* in a multigenerational context. Nevertheless, both approaches give the same geometric means and in both approaches one must use the same neutralized process (eq. 6) to tease out the effect of selection from class transmission (i.e. both use the same average reference fitness). Finally, and similarly to the above calculations of geometric means for selection, we can write

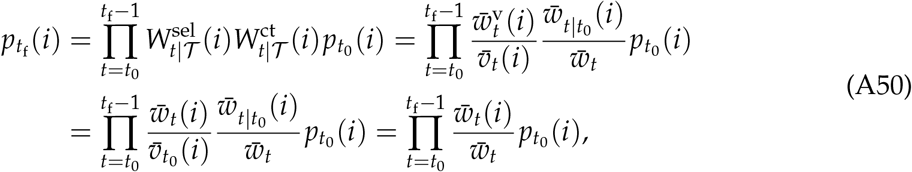

where in the third equality we used eq. (A49) and in the fourth equality eq. (A46). This calculation shows, in particular, that the contribution 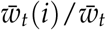 is not equal to 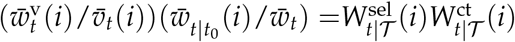 and only the latter gives the fitness describing the effect of selection and class transmission in generation *t* on the final frequency in a multigenerational context.

## A3 Asymptotic Evolutionary Process

### A3.1 Invasion Fitness

Here we derive the equations for Box B. Recall the definition of the invasion exponent of allele *i* when the initial number of such alleles is 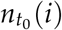:

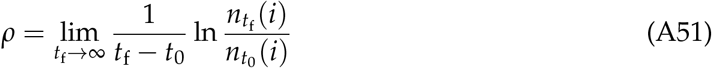

(e.g., Ferrière and Gatto, 1995, Metz et al., 1992, Tuljapurkar, 1989). Since we are interested in the growth of *i* when rare, we suppose that the model contains a steady state where *i* is absent (and while one can calculate invasion fitness for any type of population attractor, we here focus on equilibria). We thus suppose an equilibrium 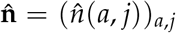 characterizing the population when *i* is absent (i.e. the ‘resident’ population) where 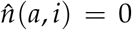 for all *a* and 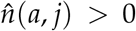 for all *j ≠ i* and all *a*, and that 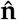 is asymptotically stable along the invariant (multidimensional) axis where allele *i* is absent (i.e. where 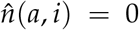 for all *a*). More specifically, we have 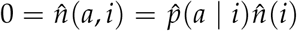 for all *a* implying that 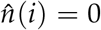. In this section and henceforth, the hat notation indicates that the quantity (state variable or a function) is evaluated at the equilibrium 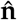.

We now calculate eq. (A51). First, from eq. (A37) we get

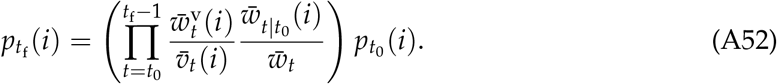

By multiplying both sides by 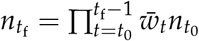 we obtain

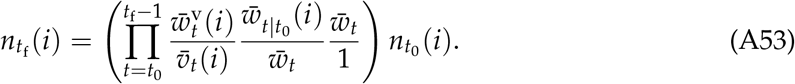

We then divide both sides by 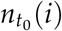 and Taylor expand the function 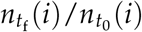 about the steady state 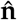 (we need to expand the whole trajectory of vectors 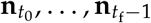), which results in the linearized equation

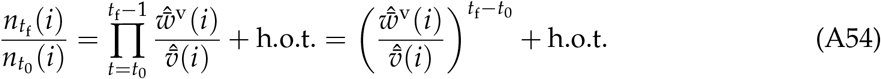

where the h.o.t. refers to terms of order 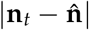 for all *t* ∈ *𝒯*. The first equality in eq. (A54) follows from evaluating 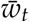 and 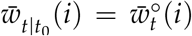 in eq. (A53) at 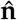: 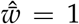 follows from the definition of an equilibrium (i.e. 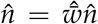 implies 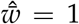), and 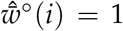 follows from the equality 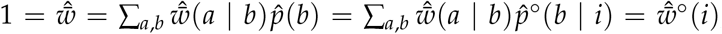, where we used the fact that at the equilibrium 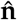 the identity 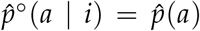 holds for all *a*. One can see this by comparing the recursion in eq. (A43) and 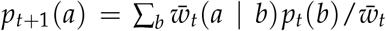, but we ought to emphasise that the identity 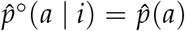 holds only for allele *i* and only at the equilibrium 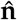 where allele *i* is absent. Furthermore, in the expressions 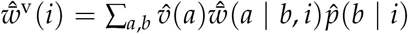 and 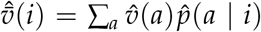 the vector 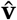 is calculated from eq. (7) and evaluated at 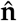 and the vector 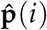 with elements 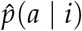 is calculated from the recursion

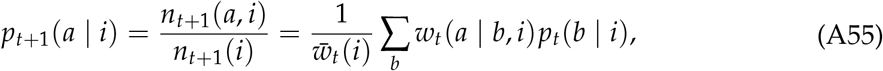

evaluated at 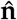 (obtained by using the definition of *p*_*t*_(*a* | *i*), eq. (1) and the recursion 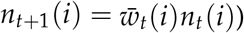). Notice that whereas 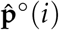 with elements 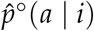 is calculated under a neutralized process from eq. (A43), the vector 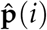 is calculated under the full evolutionary process with selection (eq. A55), and hence in general 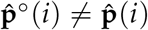.

Now, we can substitute eq. (A54) into eq. (A51) which leads to

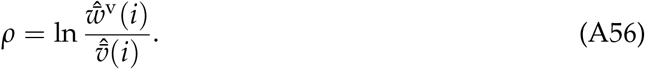

In this final step we used the fact that as *t*_f_ → ∞ the h.o.t. present in eq. (A54) converge to 0. Moreover, exp *ρ* can be shown to be the leading eigenvalue of the fitness matrix with elements 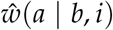 and that its corresponding eigenvector is 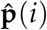. These are standard results about *ρ* and have been proved elsewhere (e.g., Caswell, 2001, Tuljapurkar, 1989) with the representation of *ρ* given in eq. (A56) appearing previously too [Lehmann and Rousset, 2020, Lehmann et al., 2016].

Finally, we show that any arbitrarily weighted fitness function could be used in eq. (A56). To this end, define an arbitrarily weighted fitness function

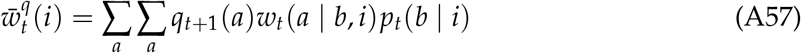

with arbitrary weights *q*_*t*_(*a*) that follow some arbitrary system of recursions. Then, we produce an analogues identity to eq. (A48), which reads as

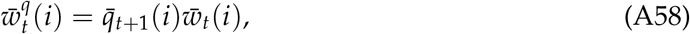

where *q*_*t*+1_(*i*) = ∑_*a*_ *q*_*t*+1_(*a*)*p*_*t*+1_(*a* | *i*). At the resident equilibrium we hence necessarily have that

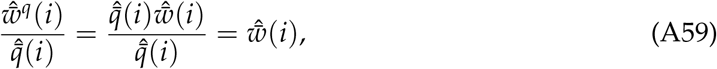

thus proving eq. (B.1) of Box B.

### A3.2 Evolutionary Dynamics under Weak Selection

Here we derive eq. (13) and show that over sufficiently large *𝒯* only the multigenerational partitioning (eqs. 8-10) is consistent with weak-selection approximations, namely, that reproductive-value weighted selection-term governs the slow phase of allele-frequency change after the initial allele-frequency change caused by class transmission has become negligible. For this, we consider models where alleles encode for closely similar phenotypes and our derivation is an adaptation to discrete time from Priklopil and Lehmann [2020, 2021] who analysed continuous-time population models.

For simplicity but without a loss of generality, let us further suppose that only two alleles segregate in the population, and that one of the two alleles, say allele *i*, is derived by mutation from the other allele, say *j*. We further suppose that the effect of the mutation on the phenotypic expression of some trait is class-specific so that the phenotypic profile of *i* can be written as **z**(*i*) = **z**(*j*) + *δ****η***, where **z**(*k*), with elements *z*(*a, k*), is the phenotypic profile of allele *k* = *i, j* such that when the allele is in class *a* it expresses phenotype *z*(*a, k*) and where ***η*** gives the direction of the effect of the mutation, with the effect in class *a* being *η*(*a*). Assuming the fitness functions to be continuous, a small parameter *δ* implies *i* and *j* having closely similar phenotypes, and hence having closely similar fitness functions. In particular, for *δ* = 0, the mutant *i* and the resident (wild-type) *j* fitness’s are equal, i.e. we have the consistency relation

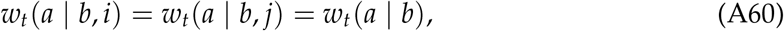

where *w*_*t*_(*a* | *b*) is a function that is independent of allele frequency (see also eqs. A5 for the notation).

Now, consider the multigenerational partitioning (eqs. 8-10) as prescribed in the main text 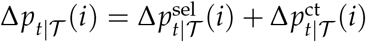 with

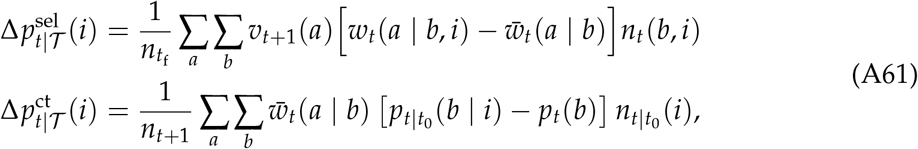

where these expressions follows from rearrangement using the definition of individual fitness functions. As we are interested in the dynamics for small values of *δ*, we Taylor expand eq. (A61) about *δ* = 0 to get

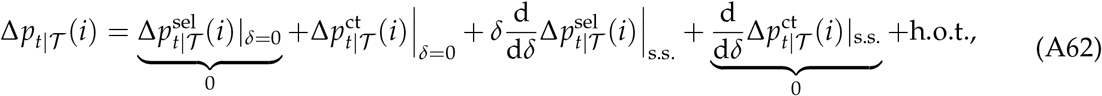

where h.o.t. refers to ‘higher order terms’ (in the above equation these terms are of order *δ*^2^) and where the first-order term is evaluated at the steady state (‘s.s.’) of the resident or wild-type population where *δ* = 0 (see below for explanation). First, we note that in eq. (A62) the zero-order term is governed purely by class transmission because the selection-term 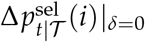 is 0 due to the fact that for *δ* = 0 we have 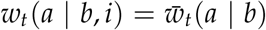 (eq. A60). The zero-order term in eq. (A62) reads

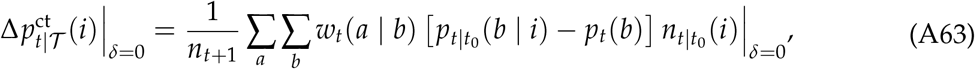

where we again used eq. (A60) and where all terms on the right-hand-side are calculated for *δ* = 0. In contrast, and second, we note that the first-order term in eq. (A62) is governed purely by selection. This is because both variables 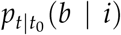 and *p*_*t*_(*b*) in eq. (A61) follow the exact same recursion 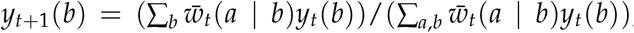, where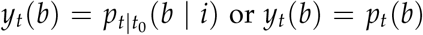, implying that (i) their derivatives are equal, and that (ii) their steady states are also equal and hence these variables will be *δ*-distance away from each other after some initial phase of evolutionary dynamics, i.e. after some finite time *t*^∗^. Because timescale-separation methods [Priklopil and Lehmann, 2020] allow us to substitute this steady state into first-order terms, we obtain that the entire derivative of class transmission 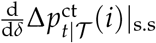 is 0. This is the reason why the first-order term in eq. (A62) is given only by the term 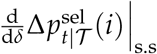 and this can further be expressed as

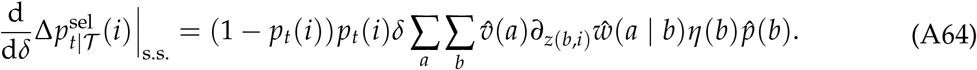

The hat notation here indicates these terms have been evaluated at the resident or wild-type steady state and ∂_*z*(*b,i*)_ refers to a partial derivative with respect to the phenotypic expression of the focal individual whose fitness function it is (calculations not shown here, see Priklopil and Lehmann 2020 for details).

Equations (A61)-(A64) thus show that the initial change in allele frequency is dominated purely by class transmission, that is, 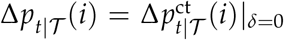. If the time of observation *𝒯* is sufficiently large so that it contains the time point *t*^∗^ after which class transmission becomes negligible, then the first-order term takes over the dynamics and at later times the allele-frequency is dominated purely by selection. That is, after *t*^∗^ the change in allele fre-quency follows 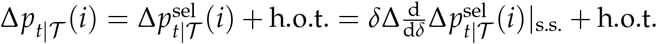. Furthermore, because any allele frequency with arbitrary weights is approximately equal to the reproductive-value weighted allele frequency and hence also 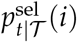 [Priklopil and Lehmann, 2021], we also have the approximation Δ*p*_*t*|*𝒯*_ (*i*) = Δ*p*_*t*_ (*i*) + h.o.t., which justifies eq. (13). Note also that the sum in eq. (A64) is a constant and defines the selection gradient for this model, and because the term in-front of the sum is sign-equivalent, the sign of the selection gradient determines whether the allele frequency increases or decreases monotonically until the allele either goes to fixation or goes extinct (Priklopil and Lehmann, 2021 for details).

Finally, we contrast the above calculation obtained from the multigenerational partitioning to that obtained from the single-generation partitioning by Taylor expanding eq. (4) about *δ* = 0 to get

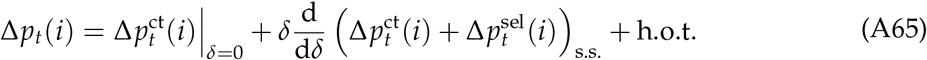

where

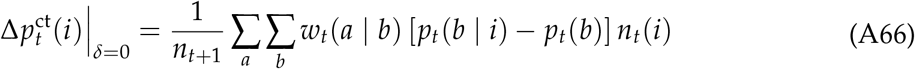

and

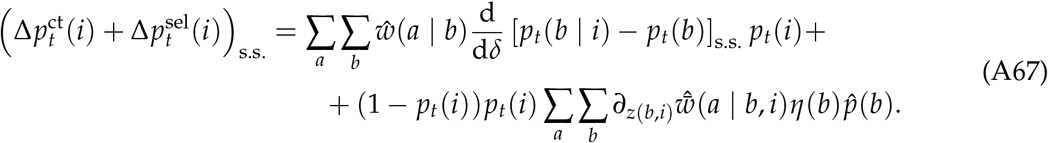

In eq. (A66) all terms on the right-hand-side are calculated for *δ* = 0. In eq. (A67), the first term on the right-hand-side does not vanish because *p*_*t*_(*b* | *i*) and *p*_*t*_(*b*) do not follow the same dynamical equation for non-zero *δ* and hence the derivative inside the brackets is nonzero, which is in contrast to 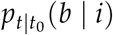 and *p*_*t*_(*b*) in eq. A63 (and note that all other derivatives are zero because they are multiplied by the term inside the brackets evaluated at *δ* = 0 which does vanish). This shows that the slow dynamics depends on the perturbation of the genetic structure (first term in eq. A67) and confirms that 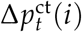 is not governed by class transmission only in the context of a multigenerational process over *𝒯* : in the single-generation partitioning one conditions on the state at time *t* and hence counts also the ancestors of individuals at *t* that underwent selection in the past [*t*_0_, *t*]. Moreover, using eqs. (A61)-(A64), we have that 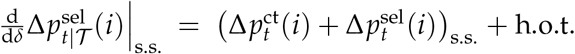 which implies that the reproductive values in the multigenerational partitioning contain the deviation term caused by genetic structure in the single-generation partitioning. While one can in principle use both eqs. (A67) and (A64) to approximate allele-frequency change under small phenotypic deviations, eq. (A64) is considerably easier to calculate. To conclude, we have confirmed that the expressions 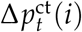 and 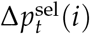 in eq. (4) do not provide a biologically satisfactory interpretation of the contributions of class transmission and selection in a multigenerational evolutionary process.

#### BOX A.

Adjoint system and the dynamics of reproductive value

Consider an arbitrary population process during *𝒯*,

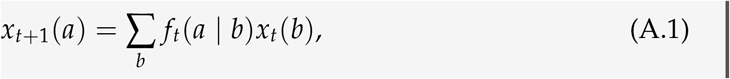

where *f*_*t*_(*a* | *b*) gives the number of type *a* individuals produced by a single type *b* individual at time *t* and *x*_*t*_(*b*) specifies the number of individuals of type *b* at time *t* under this process. In particular, this process may or may not include selection. The so-called adjoint system (e.g., Athans and Falb, 2007, p. 147) of eq. (A.1) can then be expressed as

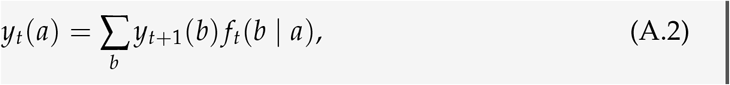

where **y**_*t*_ = (*y*_*t*_(*a*)) is called an adjoint variable (associated to the vector **x**_*t*_ = (*x*_*t*_(*a*))). A useful property of adjoint systems is that ∑_*a*_ *x*_*t*_(*a*)*y*_*t*_(*a*) is constant and equal for all *t*, and the exact value depends on the initial and final conditions 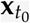 and 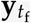. In fact, the specification of the final condition also suggests a biological interpretation for the adjoint variable **y**_*t*_. For instance, suppose that the final condition satisfies 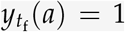 for all *a*. Then, we have 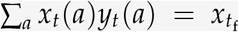 for all *t* where 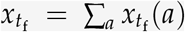 is the total population size at the final time *t*_f_ under this arbitrary population process. The adjoint variable *y*_*t*_(*a*) can be interpreted, under this arbitrary process, as the *number of individuals alive at time t*_*f*_ *that are the descendants of a single type a individual alive at time t*. Indeed, this definition follows from the recursion: the number of individuals at *t*_f_ that descend from a single individual *a* at *t* – the left hand-side of eq. (A.2) – is equal to the number *f*_*t*_(*b* | *a*) of offspring-individuals *b* that this parent individual *a* produces and where each offspring-individual at *t* + 1 is multiplied by the number *x*_*t*+1_(*b*) of its descendants alive at *t*_f_, and then we sum over all classes of offspring. Note that the final condition 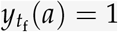 is consistent with this interpretation because the ‘descendant’ alive at *t*_f_ and its ‘ancestor’ alive at *t*_f_ must be one and the same individual, and hence the number of individuals at *t*_f_ that ‘descend’ from an individual alive at *t*_f_ must be 1. In the main text *f*_*t*_(*a* | *b*) is defined as the class-specific fitness 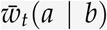, in which case eq. A.2 defines the dynamics of reproductive values under the neutralized process. Representing reproductive value dynamics as an adjoint system is also discussed e.g. in Lion [2018a, eq. 14].

#### BOX B.

Invasion fitness

Here we connect the partitioning of the recursion for a multigenerational process (eq. 5) as well as the partitioning of the geometric mean (eqs. 15-16) to the asymptotic evolutionary trajectory of a rare allele. Recall that the invasion exponent of allele *i* is defined (under some technical conditions) as 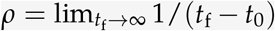 ln 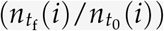 giving the asymptotic per capita rate of growth of allele *i*, which has been widely used as a summary measure of selection since at least Fisher [1930] and is often called the Malthusian growth rate. Suppose that all alleles but *i* coexist at some asymptotically stable equilibrium, which we call the resident equilibrium (the assumption of an equilibrium is not necessary as the calculations carry over to more complex attractors of the resident population e.g. Ferrière and Gatto, 1995, Metz et al., 1992, Tuljapurkar, 1989). Then, the invasion fitness for allele *i*, defined as 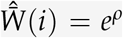, and giving the expected number of offspring produced asymptotically per generation per parent *i*, can be expressed as

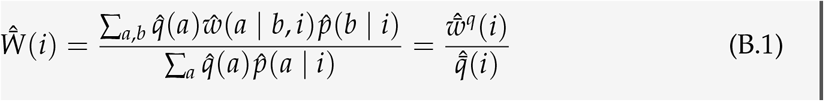

(Appendix A3.1). Here, 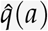 is an arbitrary weight of an individual in class *a* evaluated at the resident equilibrium (all hats hereafter indicate evaluation at an equilib-rium), 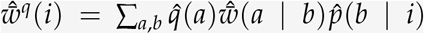 is an arbitrarily weighted fitness of *i* and 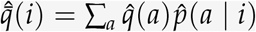. The fact that eq. (B.1) is equal for arbitrary weights implies that only selection is in play in calculations of invasion fitness. Indeed, on one hand, by setting 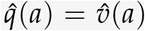 for all *a*, the invasion fitness can be represented as the asymptotic geometric mean for selection 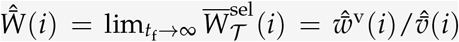 (Appendix A3.1). On the other hand, by setting 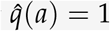 for all *a*, the invasion fitness can be represented as the asymptotic geometric mean fitness 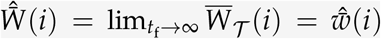 (Appendix A3.1). This confirms that the asymptotic evolutionary trajectory of a rare allele *i* is fully determined by a measure of fitness where only selection is in play. Or even more precisely, upon the introduction of the allele *i* the evolutionary trajectory is subject to both class transmission and selection but then converges to the leading eigenvector with elements 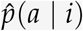, along which class transmission vanishes and the rate of growth is given by the invasion fitness determined by selection only (Appendix A3.1). We note that the two representations of invasion fitness, 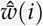 and 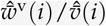, extend to spatially structured populations and 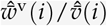 forms the basis of the inclusive fitness representation of invasion fitness [Lehmann and Rousset, 2020, Lehmann et al., 2016].

